# Nobiletin, a Polymethoxyflavonoid, Activates the Desuccinylase Activity of SIRT5 and Prevents the Development of Heart Failure

**DOI:** 10.1101/2024.01.16.575602

**Authors:** Yoichi Sunagawa, Masafumi Funamoto, Toshihide Hamabe-Horiike, Kehima Hieda, Seiichiro Yabuki, Midori Tomino, Yoshimi Ikai, Anna Suzuki, Shintaro Ogawahara, Asami Yabuta, Hana Sasaki, Ayaka Ebe, Shiomi Naito, Hidemichi Takai, Kana Shimizu, Satoshi Shimizu, Yuto Kawase, Ryuya Naruta, Yasufumi Katanasaka, Tomohiro Asakawa, Toshiyuki Kan, Kiyoshi Mori, Akira Murakami, Masahito Ogura, Nobuya Inagaki, Koji Hasegawa, Tatsuya Morimoto

## Abstract

Nobiletin is a natural compound useful for the prevention and treatment of several diseases. However, the precise role of nobiletin in heart failure is unclear. Nobiletin treatment prevents pressure overload- and myocardial infarction-induced heart failure. Using affinity purification of biotinylated nobiletin from rat heart cell lysates, we identified sirtuin 5 (SIRT5) as a novel nobiletin-binding protein. Nobiletin enhanced the desuccinylase activity of SIRT5 *in vitro*. Compared to wild-type mice, SIRT5-overexpressing transgenic mice resisted pressure overload-induced systolic dysfunction. Conversely, SIRT5 knockout disrupted the nobiletin-mediated therapeutic effects on heart failure in mice. SIRT5 desuccinylated p300 at lysine 1568 and reduced the histone acetyltransferase (HAT) activity of p300. The desuccinylated p300 mutant suppressed the phenylephrine-induced cardiomyocyte hypertrophic responses. These findings suggest that nobiletin prevents heart failure development through SIRT5-dependent inhibition of p300-HAT activity. Nobiletin, a nontoxic dietary compound, is a potential therapeutic agent for heart failure in humans.

## Introduction

Heart failure is a major clinical and public health concern worldwide^1^. Cardiac hypertrophy is usually characterized by an increase in cardiomyocyte size and left ventricular thickness in response to a pathological stress such as myocardial infarction (MI) and hypertension and is a risk factor for the development of heart failure^2,3^. Chronic stress that persists for a long period leads to irreversible cardiac hypertrophy and causes pathological cardiac remodeling and the development of heart failure associated with cardiac dysfunction. Although pharmacological strategies targeting neurohumoral factors are well-established, the five-year survival rate of patients with severe heart failure remains low^4^. Therefore, innovative pharmacotherapies are required to prevent severe heart failure.

Various neurohumoral factors bind to receptors on the surface of cardiomyocytes and activate intracellular signaling pathways. These signals eventually reach the cardiomyocyte nuclei and change the hypertrophic response gene expression pattern, resulting in cardiac hypertrophy and pathological remodeling^5^. Nuclear acetylation and deacetylation play central roles in regulating nuclear transcriptional activation during cardiac hypertrophy^5^. The intrinsic transcriptional coactivator p300 exhibits histone acetyltransferase (HAT) activity and can induce cardiac hypertrophy via the acetylation of GATA4 and MEF2^5–8^. Inhibitors of p300-HAT activity, such as curcumin, its analogs, and EPA/DHA, prevent systolic dysfunction by inhibiting nuclear acetylation^9–12^. Therefore, small-molecule compounds that regulate nuclear acetylation/deacetylation have the potential to prevent heart failure development^13^.

Natural products have potential health benefits and may be used to prevent and treat cardiovascular diseases^14,15^. Nobiletin, a polymethoxyflavonoid, is a natural compound derived from *Citrus depressa* that has various biological activities, such as anti-inflammatory, antitumor, hepatoprotective, and neuroprotective effects^16–19^. The cardioprotective effects of nobiletin have been previously reported^20–22^. Nobiletin attenuates the adverse effects of acute MI in rats^21^ and protects against pressure overload-induced cardiac hypertrophy by inhibiting NAPDH oxidase^22^. Nobiletin may also alleviate myocardial ischemia and reperfusion injury^23^. Nobiletin, a small-molecule compound, exerts pharmaceutical effects by binding to its target protein, which regulates intercellular signaling pathways and alters its activity^24,25^. However, whether nobiletin directly regulates cardiomyocyte hypertrophy and the target molecules remain unknown.

Sirtuin 5 (SIRT5) is a member of the class III histone deacetylase (HDACs) family. In contrast to the deacetylation activity of other SIRT family members, SIRT5 is a unique enzyme owing to its specific catalytic function of low deacetylation activity and high desuccinylation, demalonylation, and deglutarylation activities^26^. Although the functions of SIRT5 in the heart have been reported^27–30^, whether SIRT5 directly regulates pathological cardiac hypertrophy and the development of heart failure *in vivo* is unclear.

In this study, we demonstrate that nobiletin prevents pathological hypertrophy and systolic dysfunction in MI and transverse aortic constriction (TAC) models and identify SIRT5 as a target molecule of nobiletin. SIRT5 overexpression suppresses pressure overload-induced cardiac hypertrophy and systolic dysfunction in mice. Loss of SIRT5 disrupts the nobiletin-mediated inhibition of the development of heart failure. Moreover, SIRT5 desuccinylates p300, decreases p300-HAT activity, and negatively regulates hypertrophic responses in cardiomyocytes. Taken together, our findings demonstrate that nobiletin possesses therapeutic potency against heart failure through regulation of the SIRT5/p300 axis.

## Materials and Methods

### Methods

The Materials and Methods section is available in the Supplemental Materials and Methods.

## Results

### A polymethoxyflavonoid, nobiletin, significantly suppresses Phenylepherineinduced hypertrophic responses in cultured cardiomyocytes

To identify therapeutic compounds with antihypertrophic effects, we performed a screening assay using a natural product library^31^ and found that a low dose (150 nM) of nobiletin derived from *Citrus depressa* (Figure 1A) significantly prevented Phenylepherine (PE)-induced cardiomyocyte hypertrophy (Figures 1B-1C). Nobiletin also significantly suppressed the PE-induced hypertrophic response gene promoter activity of atrial natriuretic factor (ANF) and endothelin-1 (ET-1) (Figure 1D and Supplemental Figure S1).

**Figure.1.**
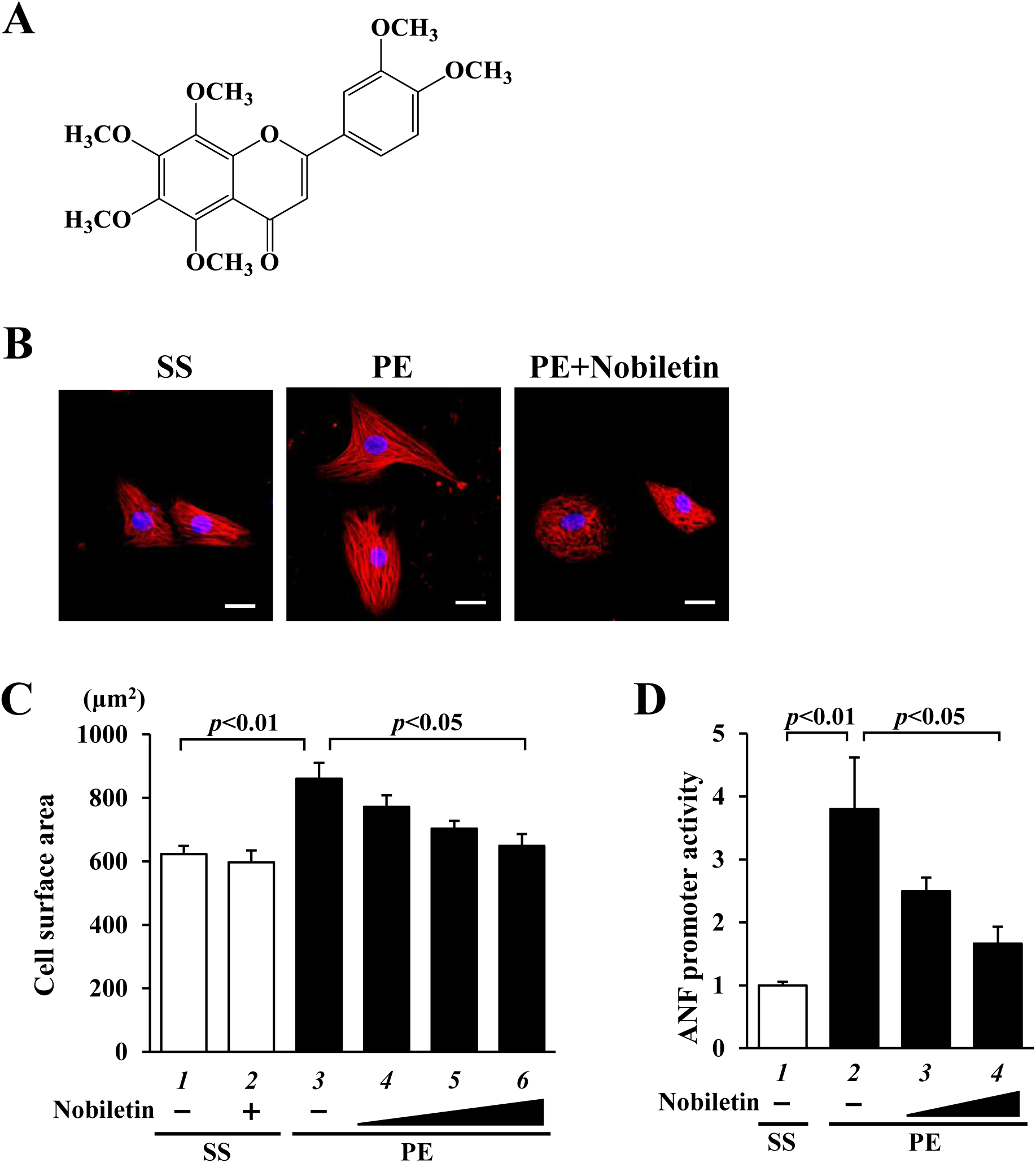
A natural polymethoxy flavonoid, nobiletin significantly prevents PE-induced hypertrophic responses in cultured cardiomyocytes. **(A)** The structure of Nobiletin. **(B)** Cultured cardiomyocytes prepared from neonatal rats were treated with nobiletin (15, 50, 150 nM) for 2 h and then stimulated with 30 μM PE for 48 h. The representative images of cardiomyocytes were stained with anti-β-MHC antibody (red) and Hoechst33258 (blue). Scar bar indicates 20 μm. **(C)** The cell surface area of β-MHC-positive cardiomyocyte was measured by Image J. The values were shown as means ± SEM. **(D)** Cardiomyocytes were transfected with pANF-luc and pRL-SV40 in the presence or absence of nobiletin for 2 h and then stimulated with PE for 48 h. The relative ANF promoter activity was calculated from the ratio of firely luciferase activity to sea pansy luciferase activity. The values were shown as means ± SEM from three independent experiments, and each was carried out in duplicate.

These results suggest that nobiletin suppresses PE-induced hypertrophic responses in cultured cardiomyocytes.

### Nobiletin significantly suppresses TAC-induced heart failure development in vivo

To determine whether nobiletin prevents *in vivo* heart failure development, we used two animal models of heart failure: TAC mice and MI rats. One day after surgery, C57BL6j mice subjected to TAC or sham surgery were orally administered nobiletin (20 mg/kg/day) or vehicle for eight weeks. The echocardiographic results showed that nobiletin treatment significantly suppressed TAC-induced cardiac hypertrophy and systolic dysfunction in mice (Figures 2A-2C and Supplemental Table S2). As indicated by the heart weight per body weight (HW/BW), cardiac hypertrophy was significantly inhibited by nobiletin treatment (Figure 2D and Supplemental Figure S2A). Nobiletin also prevented the TAC-induced increases in ANF and brain natriuretic peptide (BNP) mRNA levels (Figures 2E-2F). Hematoxylin and eosin (HE) and Masson’s trichome (MT) staining showed that TAC-induced myocardial cell hypertrophy and perivascular fibrosis were significantly suppressed by nobiletin treatment (Figures 2G-2J).

**Figure. 2.**
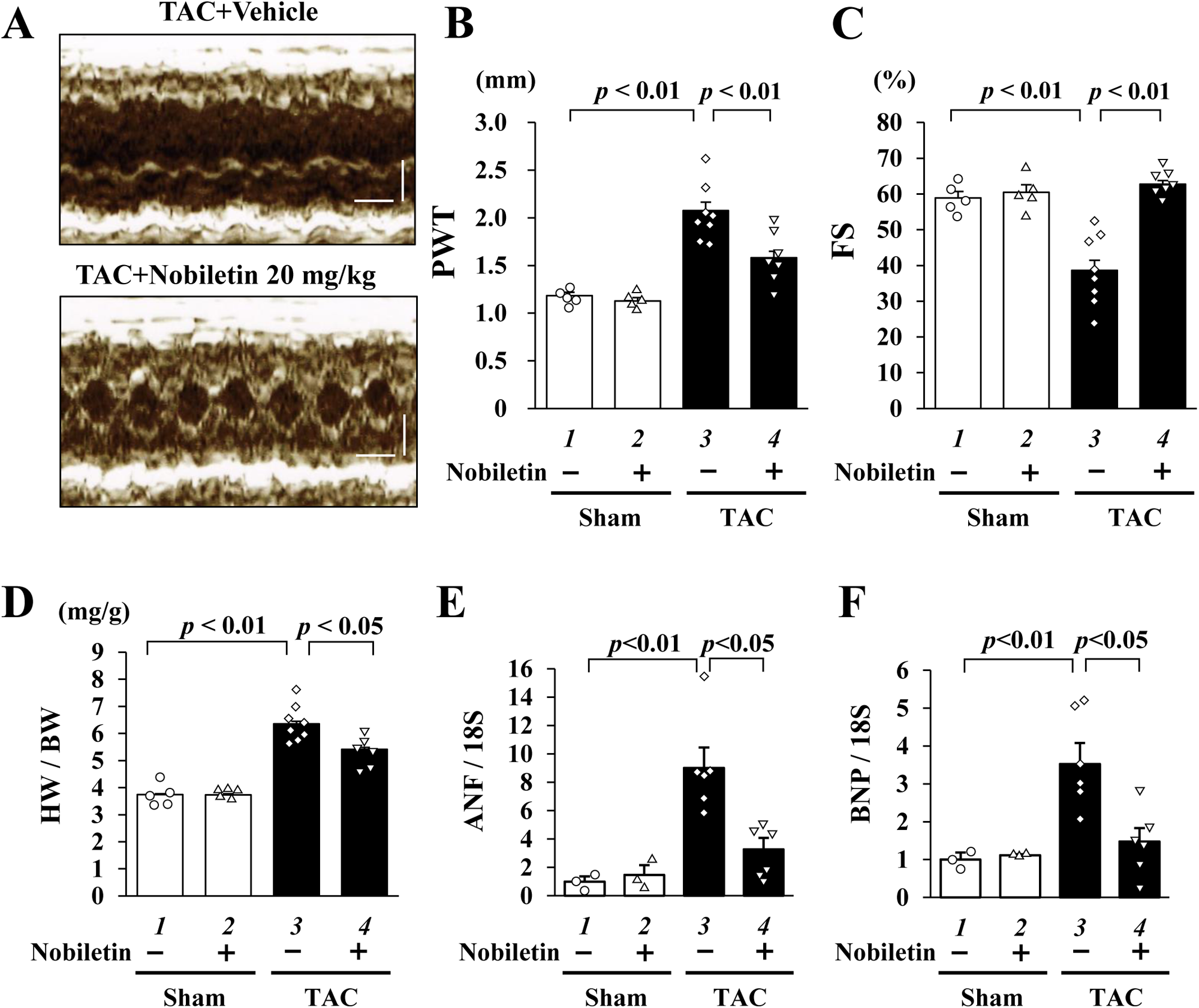
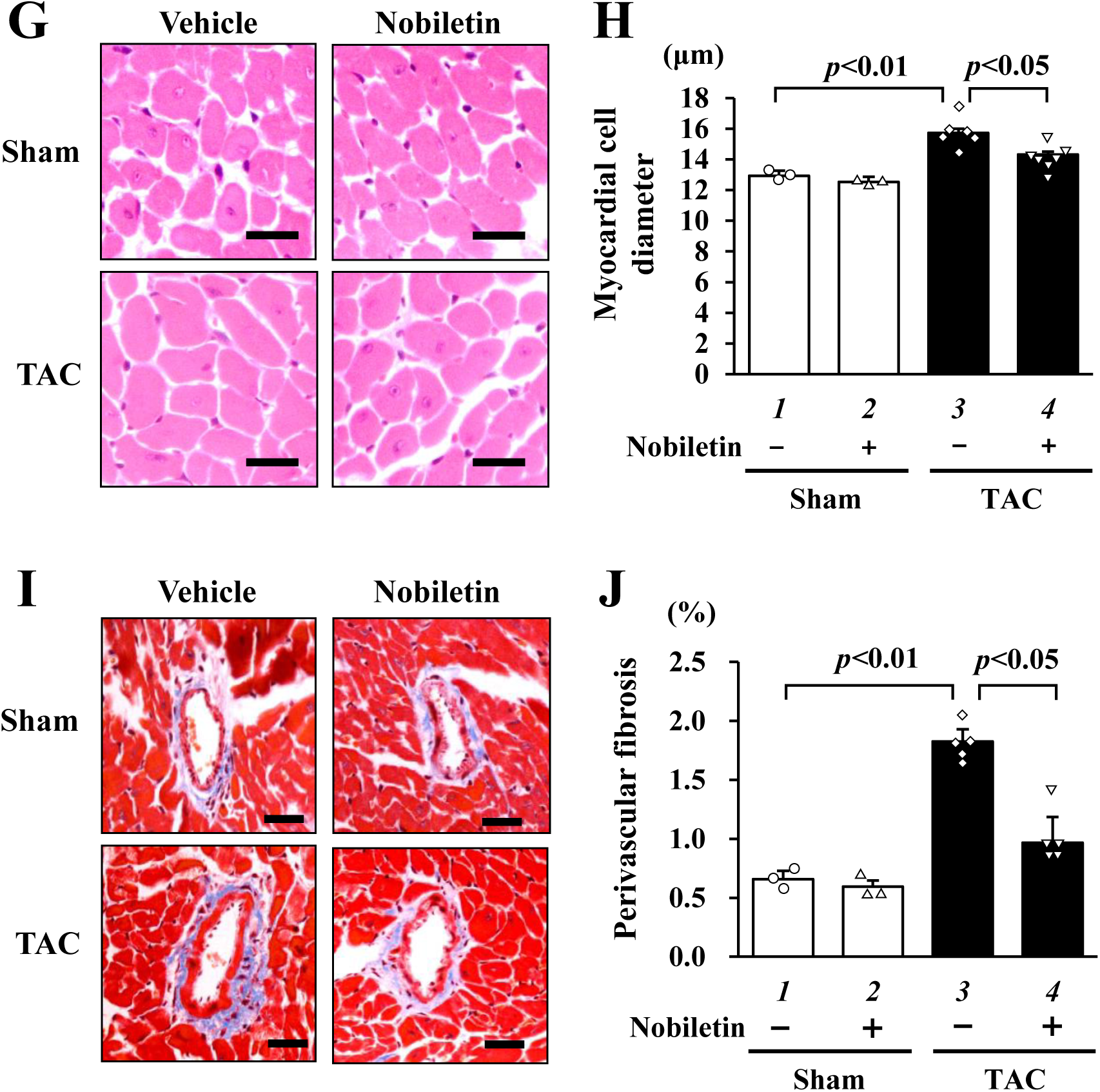
Nobiletin significantly prevents pressure overload-induced development of heart failure in mice. One day after surgery, mice with TAC or sham mice were treated with nobiletin (20 mg/kg/day) or vehicle (1% gum arabic) for 8 weeks. **(A)** The representative M-mode echocardiography images were from indicated groups. The vertical line indicates 2 mm and the horizontal line indicates 0.2 sec. **(B and C)** PWT **(B)** and FS **(C)** values were shown as means ± SEM. Other parameters were shown in Supplemental Table S2. **(D)** The ratio of HW/BW was shown as means ± SEM (mg/g). **(E and F)** Total RNA from each heart was subjected to quantified RT-PCR. The mRNA levels of ANF **(E)** and BNP **(F)** were normalized by 18S. **(G)** The representative images of cross-sectional myocardial cells were stained with HE. Scar bar: 10 μm. **(H)** The area of myocardial cell diameter was measured for at least 50 cells in each heart. **(I)** The representative images of perivascular fibrosis were stained with MT in each group. Scar bar: 20 μm. **(J)** The area of perivascular fibrosis was measured for at least five intramyocardial coronary arteries with a lumen size over 20 μm in each group. The values were shown as mean ± SEM.

One week after MI surgery, Sprague Dawley (SD) rats with moderate MI (fractional shortening [FS]<40%) were orally administered low- or high-dose (2 or 20 mg/kg/day, respectively) nobiletin or vehicle for six weeks. Before treatment, echocardiographic and hemodynamic parameters did not differ among the groups (Supplemental Table S3). Six weeks after treatment, a high dose of nobiletin significantly suppressed the MI-induced increase in the posterior wall thickness (PWT) and decreased FS, a parameter of systolic function (Figures 3A-3C and Supplemental Table S4). MI-induced upregulation of ANF and BNP mRNA levels was significantly prevented by nobiletin treatment (Figures 3D-3E). Histological analysis revealed that a high dose of nobiletin significantly suppressed MI-induced increases in the myocardial cell diameter and perivascular fibrosis (Figures 3F-3I). The infarct areas did not differ between the MI groups (Supplemental Figure S3 and Table S4). These results suggest that nobiletin prevents heart failure development in the two different chronic heart failure models.

**Figure. 3.**
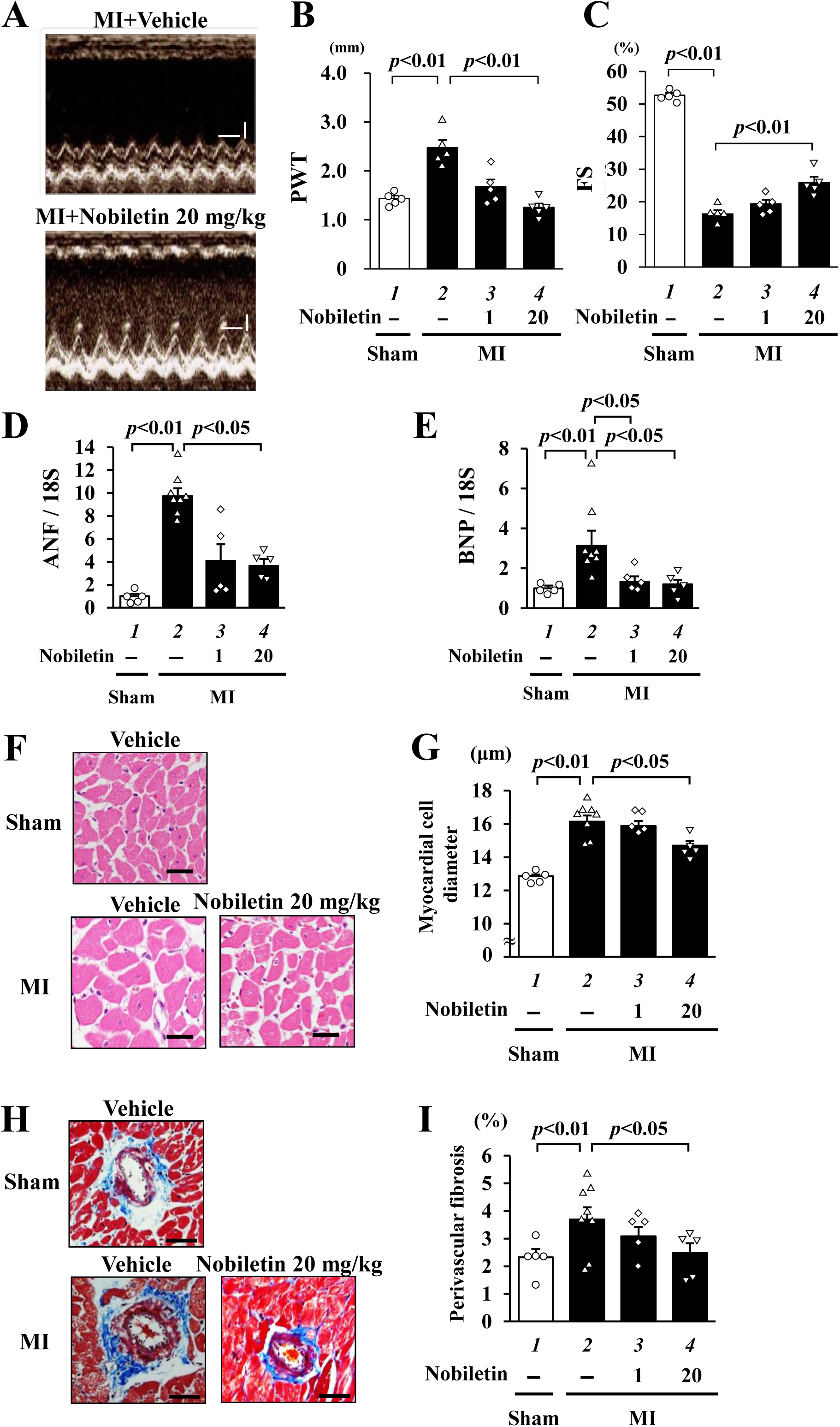
Nobiletin significantly prevents MI-induced development of heart failure in rats. At one week after surgery, rats with moderated MI (FS<40%) were treated with nobiletin (1 and 20 mg/kg/day) or vehicle (1% gum arabic) for 6 weeks. **(A)** The representative images of M-mode echocardiography from indicated groups at seven weeks after surgery. The vertical line indicates 2 mm and the horizontal line indicates 0.2 sec. **(B and C)** PWT **(B)** and FS **(C)** values were shown as means ± SEM. Other parameters were shown in Supplemental Tables S2 and S3. **(D and E)** Total RNA from each heart was subjected to quantified RT-PCR. The mRNA levels of ANF **(D)** and BNP **(E)** were normalized by 18S. **(F)** The representative images of cross-sectional myocardial cells were stained with HE. Scar bar: 20 μm. **(G)** The area of myocardial cell diameter was measured for at least 50 cells in each heart. **(H)** The representative images of perivascular fibrosis were stained with MT in each group. Scar bar: 40 μm. **(I)** The area of perivascular fibrosis was measured for at least five intramyocardial coronary arteries with a lumen size > 20 μm in each group. The values were shown as mean ± SEM.

### SIRT5 is a novel target molecule of nobiletin

To determine the molecular target of nobiletin for hypertrophic responses, we synthesized nobiletin derivatives using a 5-position carbon probe conjugated with biotin (Bio-Nobi, Supplemental Figure S4A) or TokyoGreen, a fluorescent substrate (TG-Nobi,^2^ Supplemental Figure S5A)^32^. Cell surface area measurements revealed that the antihypertrophic effect of Bio-Nobi was conserved (Supplemental Figure S4B). TG-Nobi fluorescence was observed in the nucleus and cytosol of cardiomyocytes (Supplemental Figure S5B). To purify the nobiletin-binding proteins, 30 mg of cell lysates from rat hearts were incubated with 4 µM Bio-Nobi or 4 µM biotin as a control and precipitated with streptavidin sepharose. Nobiletin-binding proteins were subjected to silver staining (Supplemental Figure S6A). Using LC-MS/MS, binding proteins were analyzed by Professor Steven P. Gygi (Harvard Medical School, USA)^33^. The LC-MS/MS results identified 162 proteins as nobiletin target candidates (Supplemental Figure S6B and Supplemental Data). Among these, we focused on SIRT5, a member of the sirtuin family^34^. To confirm the direct interaction between nobiletin and SIRT5, a binding assay was performed using Bio-Nobi and recombinant SIRT5. The results showed that Bio-Nobi was physically bound to SIRT5, and this binding was competitively disrupted with the addition of 40 µM nobiletin (Figure 4A). To clarify whether nobiletin regulates SIRT5 enzyme activity, the desuccinylase activity of SIRT5 was measured using nobiletin, resveratrol (an SIRT1 activator), and dimethyl sulfoxide (DMSO) as a control. The results showed that nobiletin, but not resveratrol, enhanced the SIRT5-mediated dessuccinylase activity *in vitro* (Figure 4B). Conversely, nobiletin decreased the SIRT5-mediated deacetylase activity (Supplemental Figure S6C). Immunofluorescence assays revealed that SIRT5 was mainly distributed in the nuclei of cardiomyocytes (Figure 4C).

**Figure. 4.**
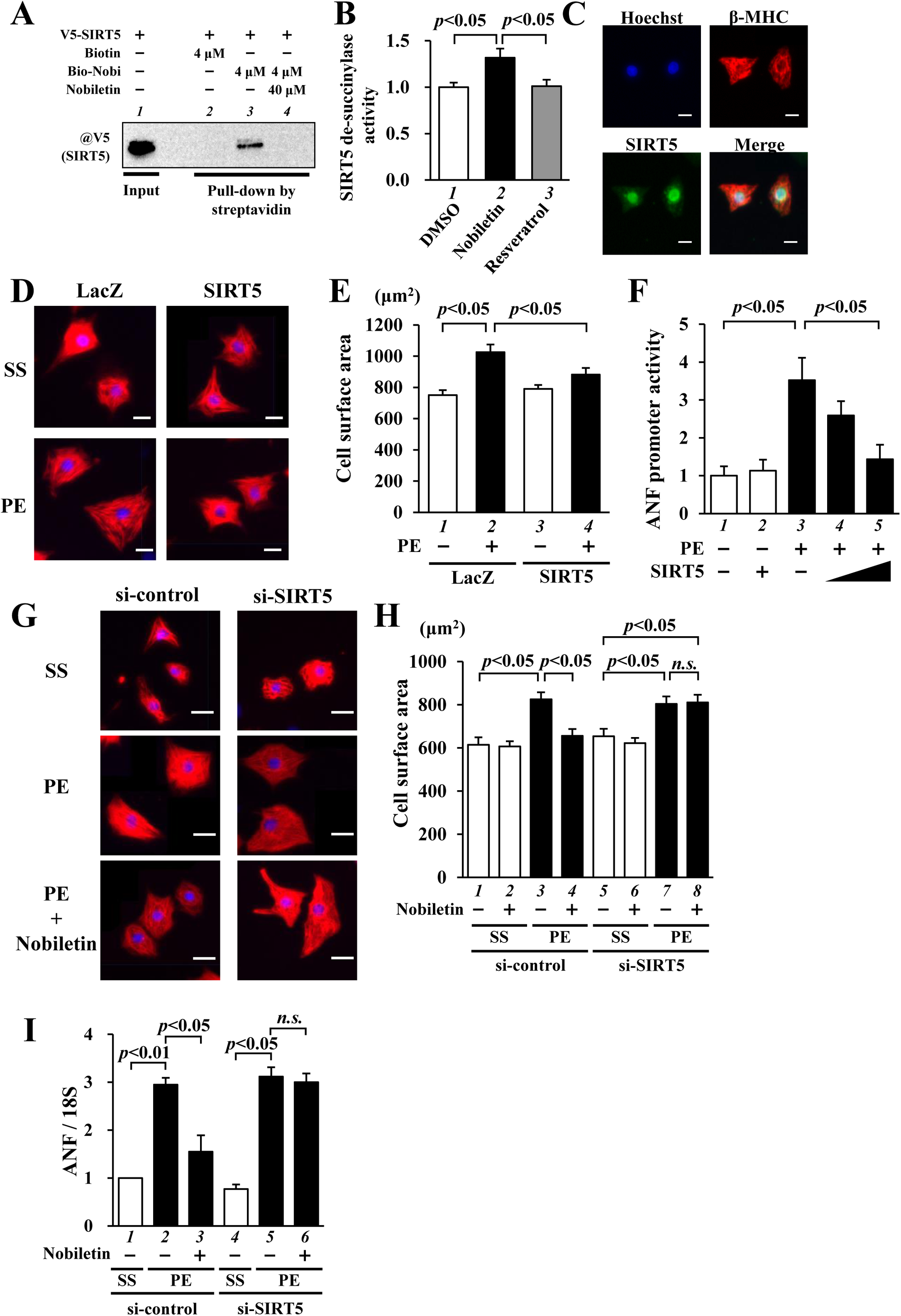
SIRT5, a target molecule of nobiletin, is required for the anti-hypertrophic effect of nobiletin. **(A)** Recombinant SIRT5-V5 prepared from *E. coli* was incubated with bio-Nobi (4 μM) alone or biotin (4 μM) in the presence or absence of nobiletin (40 μM). The binding protein was detected by western blotting using an anti-V5 antibody. **(B)** The de-succinylation activities of SIRT5 in nobiletin (100 µM) or resveratrol (100 µM) were measured using a commercial kit. The relative values were shown as means ± SEM from three independent experiments. **(C)** Cultured cardiomyocytes were stained with anti-β-MHC (red) and anti-SIRT5 (green) antibodies and Hoechst 33258 (blue). Scar bar: 20 μm. **(D)** The representative images of cardiomyocytes overexpressed using adenovirus encoding V5-tagged SIRT5 or LacZ as a control in the presence or absence of PE for 48 h. These cells were stained with anti-β-MHC (red) antibody and Hoechst33258 (blue). Scar bar: 20 μm. **(G)** The representative images of cardiomyocytes transfected with si-SIRT5 or si-control, pre-treated with nobiletin or DMSO, and then stimulated with PE for 48 h. These cells were stained with anti-β-MHC (red) antibody and Hoechst33258 (blue). Scar bar: 20 μm. **(E and H)** The cell surface area of β-MHC-positive cardiomyocytes was measured. The data were shown as means ± SEM from three independent experiments. **(F)** Cardiomyocytes were transfected with pANF-luc and pRL-SV40 in the presence or absence of pcDNA-SIRT5 and stimulated with PE for 48 h. The relative promoter activities were shown as means ± SEM from three independent experiments, and each was carried out in duplicate. **(I)** Total RNAs from indicated cardiomyocytes were subjected to quantified RT-PCR. The mRNA level of ANF was normalized by that of 18S. The data were shown as means ± SEM from three independent experiments.

To investigate the expression of SIRT5 during heart failure development, western blotting was performed using TAC- and MI-induced chronic heart failure hearts. Cardiac SIRT5 mRNA levels were reduced in TAC and MI models (Supplemental Figures S7A and S7B). According to these results, succinylated proteins in the nucleus were increased in hearts treated with TAC and MI (Supplemental Figures S7C and S7D). Therefore, SIRT5 desuccinylation activity may be involved in the development of heart failure *in vivo*. To determine whether SIRT5 regulates hypertrophic responses, we investigated SIRT5 gain or loss of function in cultured cardiomyocytes. Adenoviral overexpression of SIRT5 significantly suppressed PE-induced cardiomyocyte hypertrophy (Figures 4D-4E and Supplemental Figure S8A). A reporter gene assay showed that SIRT5 significantly decreased the PE-induced ANF and ET-1 promoter activation (Figure 4F and Supplemental Figure S8B). Although SIRT5 knockdown had no effect on PE-induced cardiomyocyte hypertrophy or hypertrophic response marker transcription, nobiletin-induced inhibition of hypertrophic responses significantly disappeared with SIRT5 knockdown (Figures 4G-4I and Supplemental Figures S8C-S8D). These results indicate that SIRT5, a nobiletin-binding protein, is essential for the antihypertrophic effects of nobiletin.

### SIRT5 overexpression significantly suppresses TAC-induced heart failure development in vivo

To determine the role of SIRT5 in the development of heart failure, transgenic mice overexpressing SIRT5 driven by the CAG promoter (SIRT5-TG mice) were used^35^. The cardiac expression level of SIRT5 was 5.9-fold higher in SIRT5-TG mice than in their WT littermates (Supplemental Figures S9A-S9B). SIRT5-TG and WT mice exhibited similar echocardiographic and hemodynamic parameters under normal conditions. Eight-week-old SIRT5-TG and WT mice were subjected to TAC or sham surgery. Eight weeks after surgery, echocardiographic analysis was performed. SIRT5 overexpression significantly prevented the TAC-induced PWT increases and FS decreases (Figures 5A-5C and Supplemental Table S5). Moreover, TAC-induced increases in HW/BW and ANF, BNP, and β-MHC mRNA levels were significantly inhibited in SIRT-TG mice, as compared to those in WT mice (Figures 5D-5F and Supplemental Figures S9C-S9D). Myocardial cell hypertrophy and perivascular fibrosis were significantly suppressed in SIRT5-TG mice (Figures 5G-5J). These results indicate that SIRT5 overexpression suppresses pressure overload-induced cardiac hypertrophy and systolic dysfunction in mice.

**Figure. 5.**
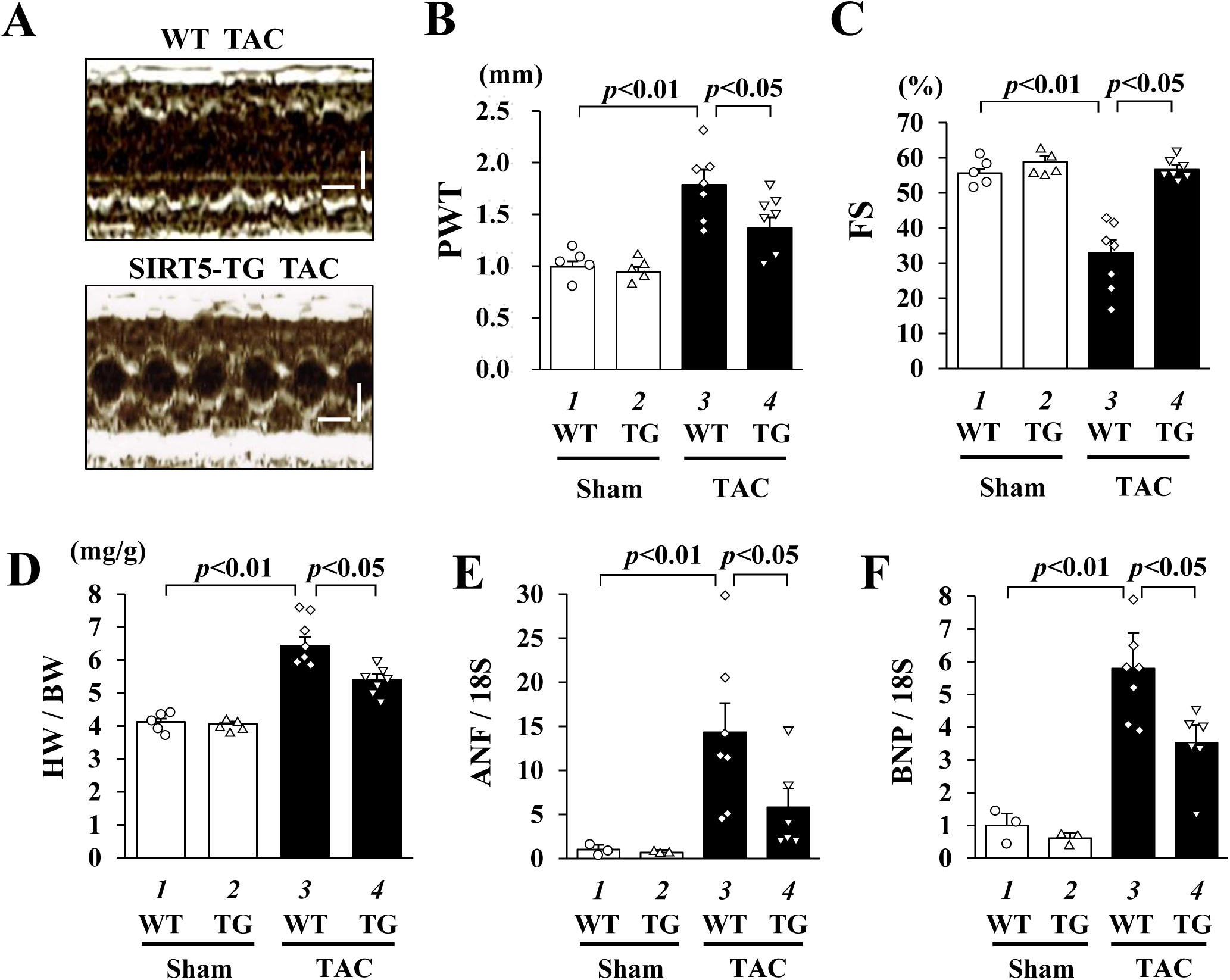
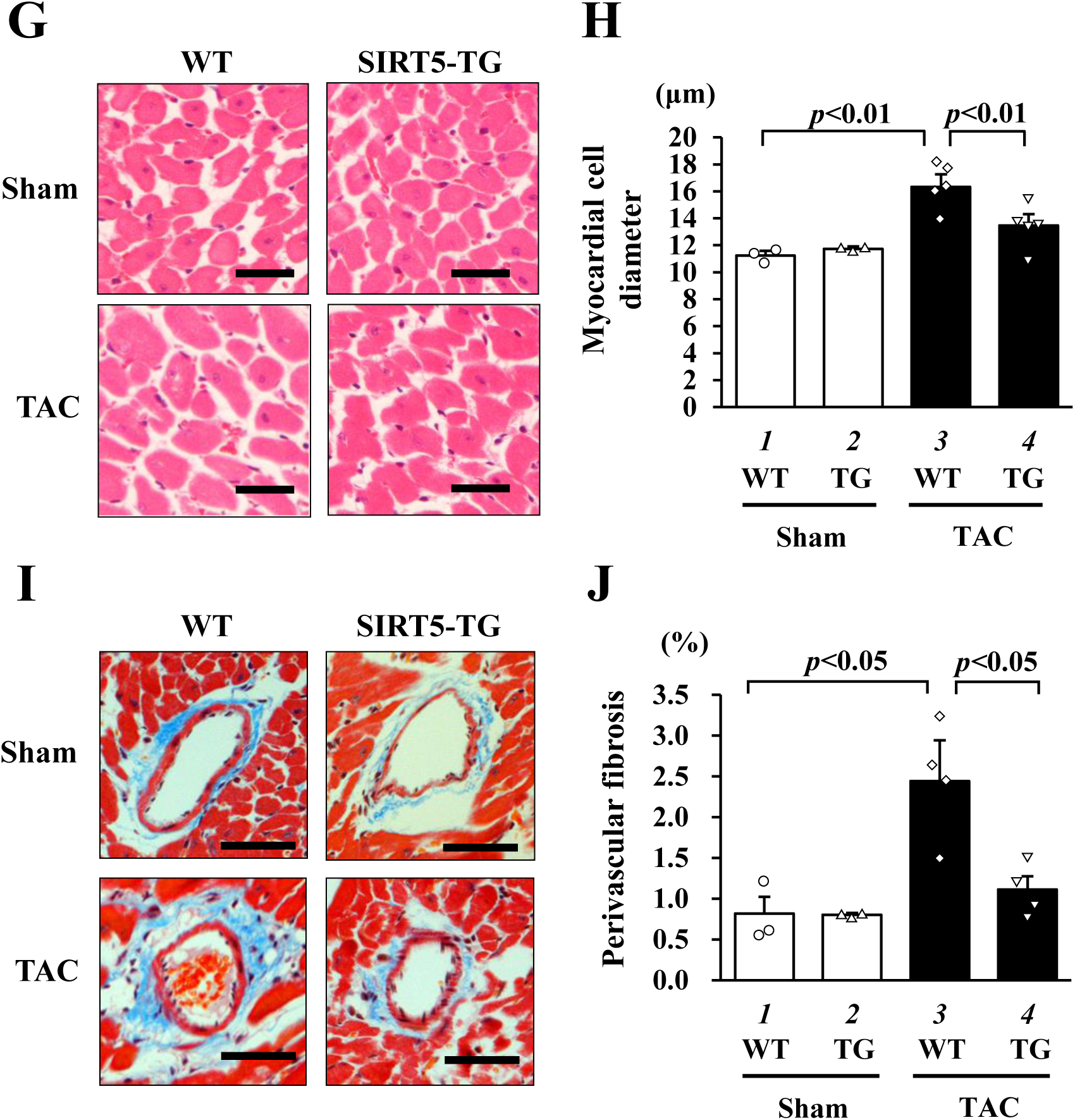
Overexpression of SIRT5 significantly attenuated pressure overload-induced development of heart failure in mice. SIRT5-TG and WT mice were subjected to TAC or sham surgery. **(A)** The representative images of M-mode echocardiography from indicated groups at eight weeks after surgery. **(B and C)** The values of PWT **(B)** and FS **(C)** were shown as means ± SEM. Other parameters were shown in Supplemental Table S5. **(D)** The ratio of HW/BW was shown as means ± SEM (mg/g). **(E and F)** Total RNA from each heart was subjected to quantified RT-PCR. The mRNA levels of ANF **(E)** and BNP **(F)** were normalized by 18S. **(G)** The representative images of cross-sectional myocardial cells were stained with HE. Scar bar: 20 μm. **(H)** The area of myocardial cell diameter was measured for at least 50 cells in each heart. **(I)** The representative images of perivascular fibrosis were stained with MT in each group. Scar bar: 20 μm. **(J)** The area of perivascular fibrosis was measured for at least five intramyocardial coronary arteries with a lumen size > 20 μm in each group. The values were shown as mean ± SEM.

### SIRT5 is essential for the therapeutic effect of nobiletin on TAC-induced heart failure

To investigate whether SIRT5 is involved in the therapeutic effects of nobiletin *in vivo*, we used SIRT5 knockout mice (SIRT5-KO mice) (Supplemental Figures S10A-S10B). The level of succinylated cardiac proteins in the nucleus was higher in homogeneous SIRT5-KO mice than in heterogeneous SIRT5-KO and WT littermate mice (Supplemental Figure S10C). Homogenous SIRT5-KO and WT mice were subjected to TAC or sham surgery and then orally administered daily nobiletin (20 mg/kg/day) or vehicle one day after surgery. Although nobiletin significantly improved TAC-induced cardiac hypertrophy and systolic dysfunction in WT mice, the therapeutic effects of nobiletin disappeared in SIRT5-KO mice (Figures 6A-6C and Supplemental Table S6). The cardiac weight and size were similar between the nobiletin and vehicle groups in SIRT5-KO mice that had undergone TAC surgery (Figure 6D and Supplemental Figures S10D). The nobiletin-induced inhibition of TAC-induced increases in ANF and BNP mRNA levels was lost in SIRT5-KO mice, as compared to that in WT mice (Figure 6E and Supplemental Figure S10E). Histological analysis showed that nobiletin treatment prevented TAC-induced increases in myocardial cell hypertrophy and perivascular fibrosis in WT mice but not in SIRT5-KO mice (Figures 6F-6I). These results suggest that SIRT5 is required for the therapeutic effects of nobiletin on cardiac hypertrophy and heart failure development in mice.

**Figure. 6.**
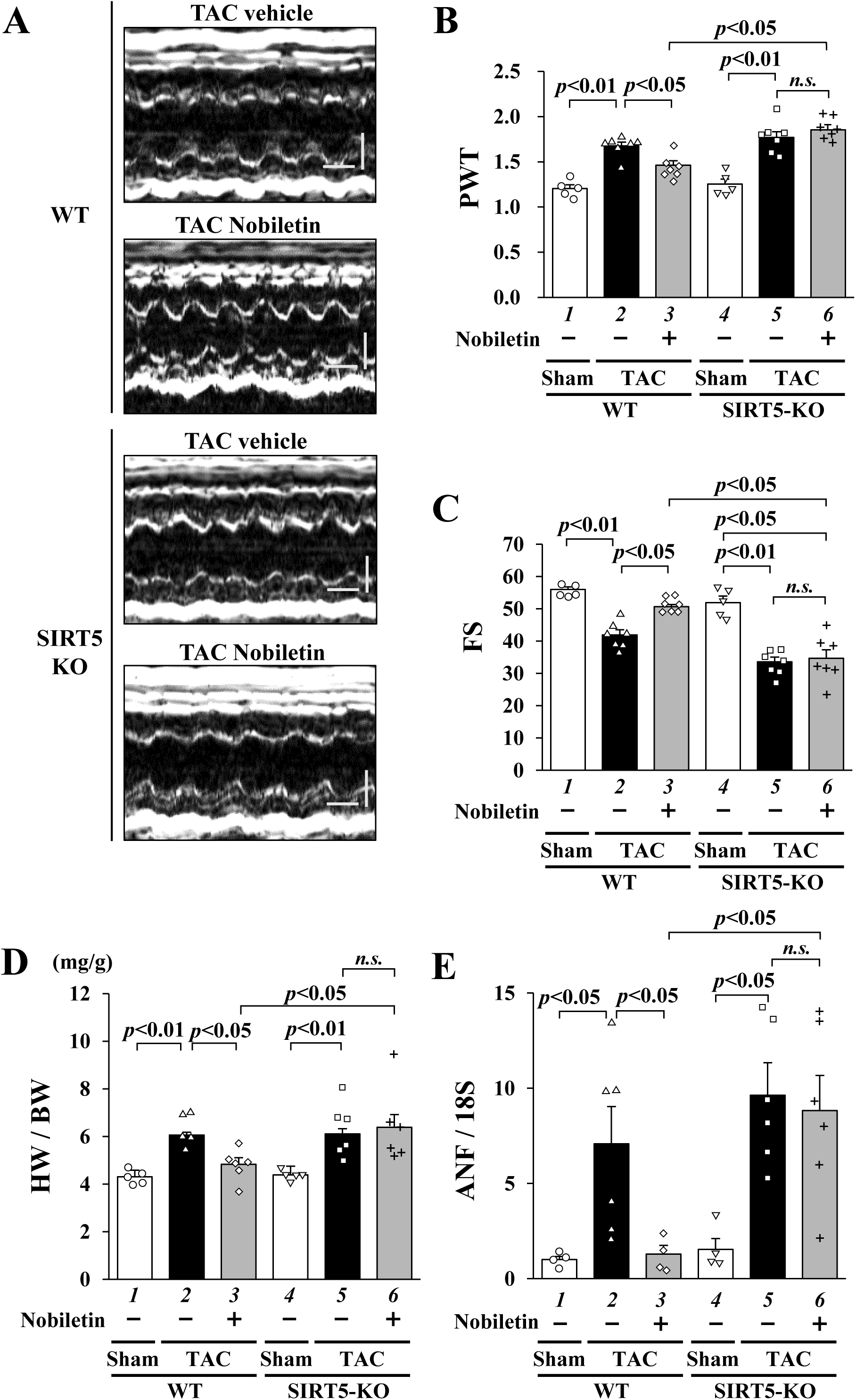
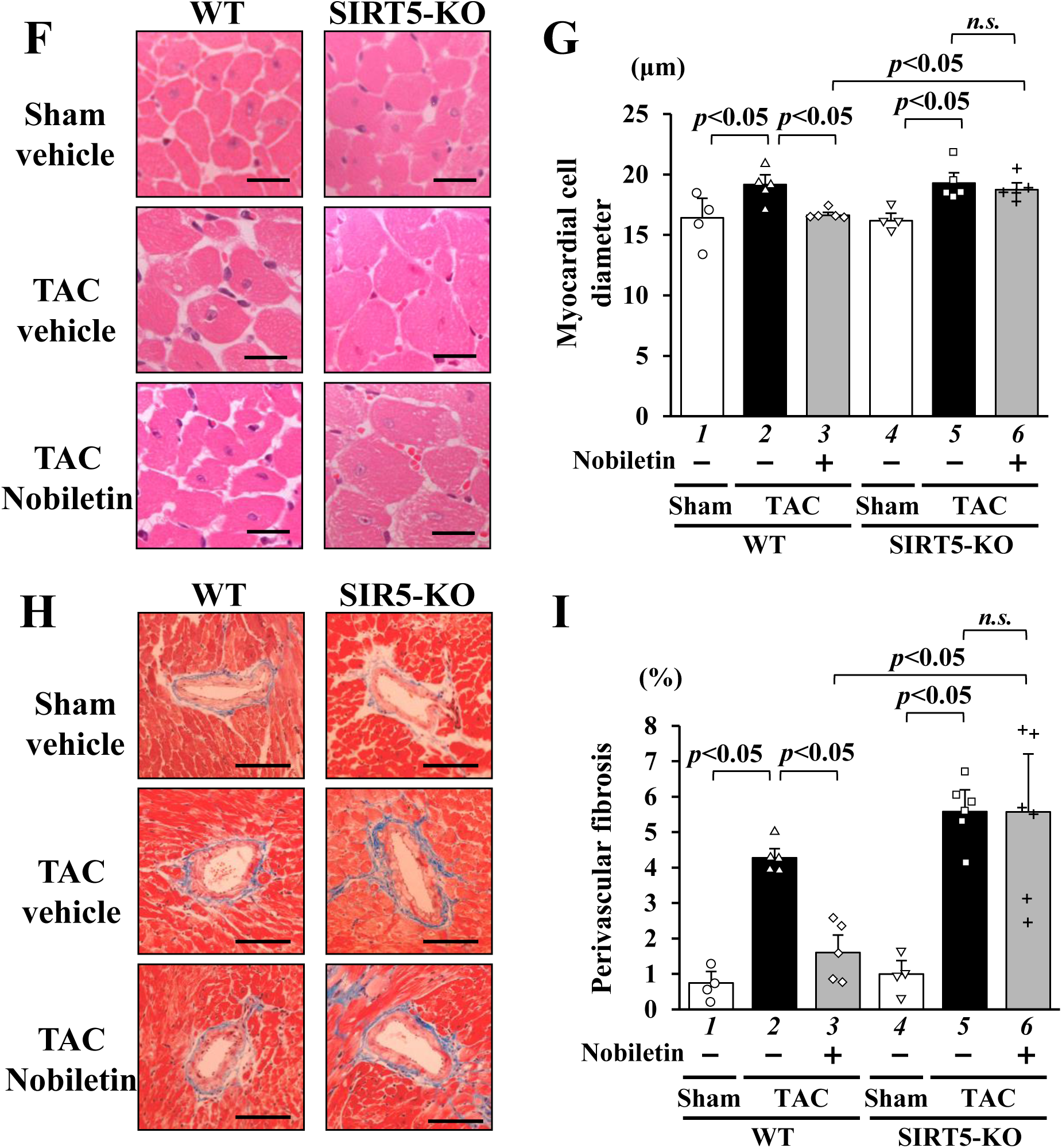
SIRT5 is required for the therapeutic potency of nobiletin on the development of heart failure in mice. SIRT5-KO and WT mice were subjected to TAC or sham surgery. At one day after surgery, mice were orally treated with nobiletin (20 mg/kg/day) or vehicle (1% gum arabic) for 6 weeks. **(A)** The representative M-mode echocardiography images from indicated groups six weeks after surgery. **(B and C)** PWT **(B)** and FS **(C)** values were indicated as means ± SEM. Other parameters were shown in Supplemental Table S6. **(D)** The ratio of HW/BW was shown as means ± SEM (mg/g). **(E)** Total RNA from each heart was subjected to quantified RT-PCR. The mRNA level of ANF was normalized by that of 18S. **(F)** The representative images of cross-sectional myocardial cells were stained with HE. Scar bar: 20 μm. **(G)** The area of myocardial cell diameter was measured for at least 50 cells in each heart. **(H)** The representative images of perivascular fibrosis were stained with MT in each group. Scar bar: 40 μm. **(I)** The area of perivascular fibrosis was measured for at least five intramyocardial coronary arteries with a lumen size > 20 μm in each group. The values were shown as mean ± SEM.

### SIRT5 desuccinylates p300 and decreases p300-HAT activity and hypertrophic responses

Since both SIRT5 overexpression and nobiletin treatment prevented the PE-induced promoter activities of ANF and ET-1, which are controlled by the p300/GATA4 pathway^36,37^, nobiletin/SIRT5 may regulate the p300/GATA4 pathway^36,37^. To test this hypothesis, we investigated the effect of SIRT5 on the p300-induced acetylation of GATA4 and promoter activation in HEK293T cells. Immunoprecipitation western blotting revealed that SIRT5 overexpression significantly suppressed p300-induced GATA4 acetylation without affecting the expression level of p300 or binding between p300 and GATA4 (Figures 7A-7C). Moreover, a reporter assay showed that p300/GATA4-dependent activation of ANF and ET-1 transcription was significantly inhibited in a SIRT5 expression level-dependent manner (Figures 7D and Supplemental Figure S11A).

**Figure. 7.**
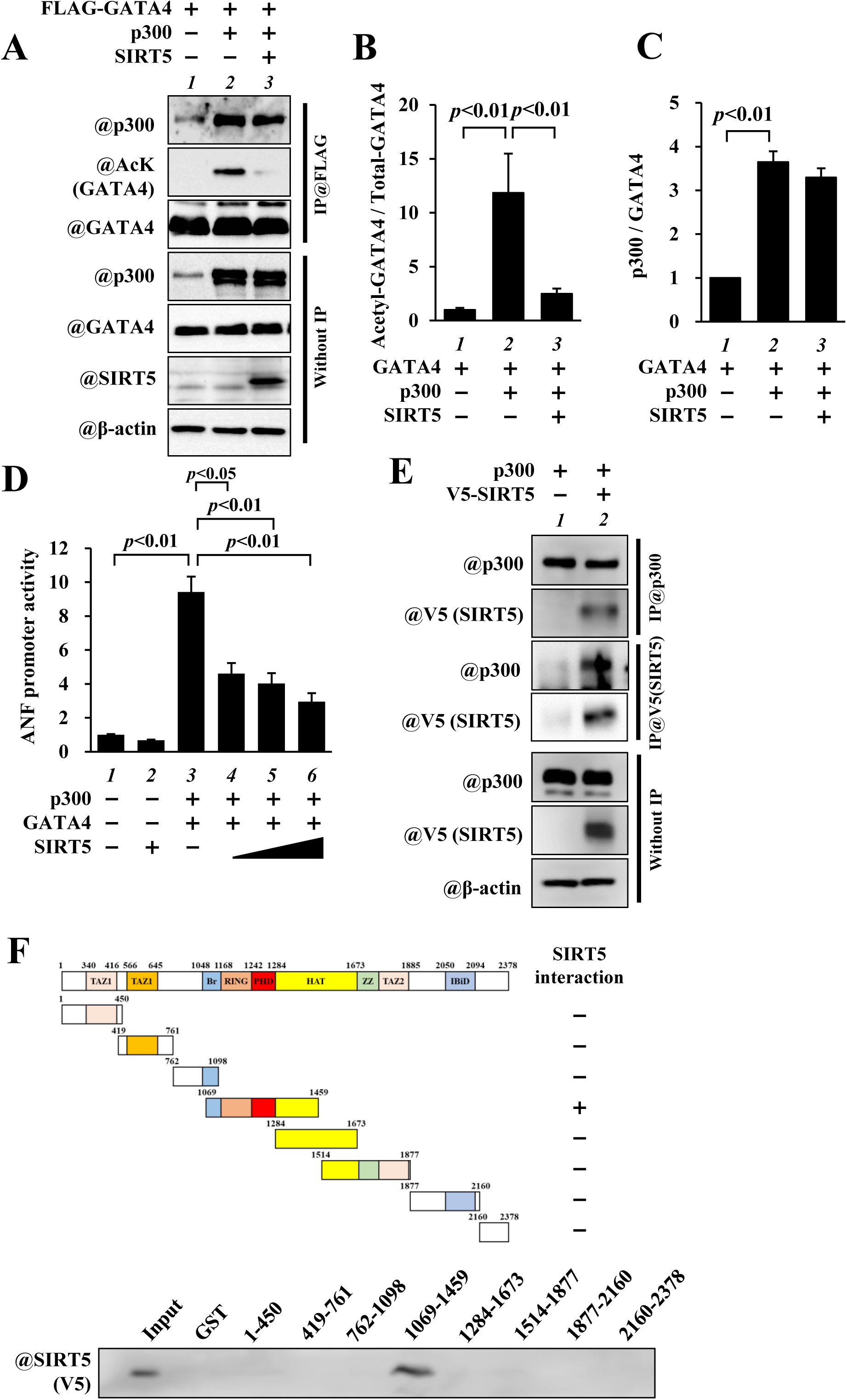
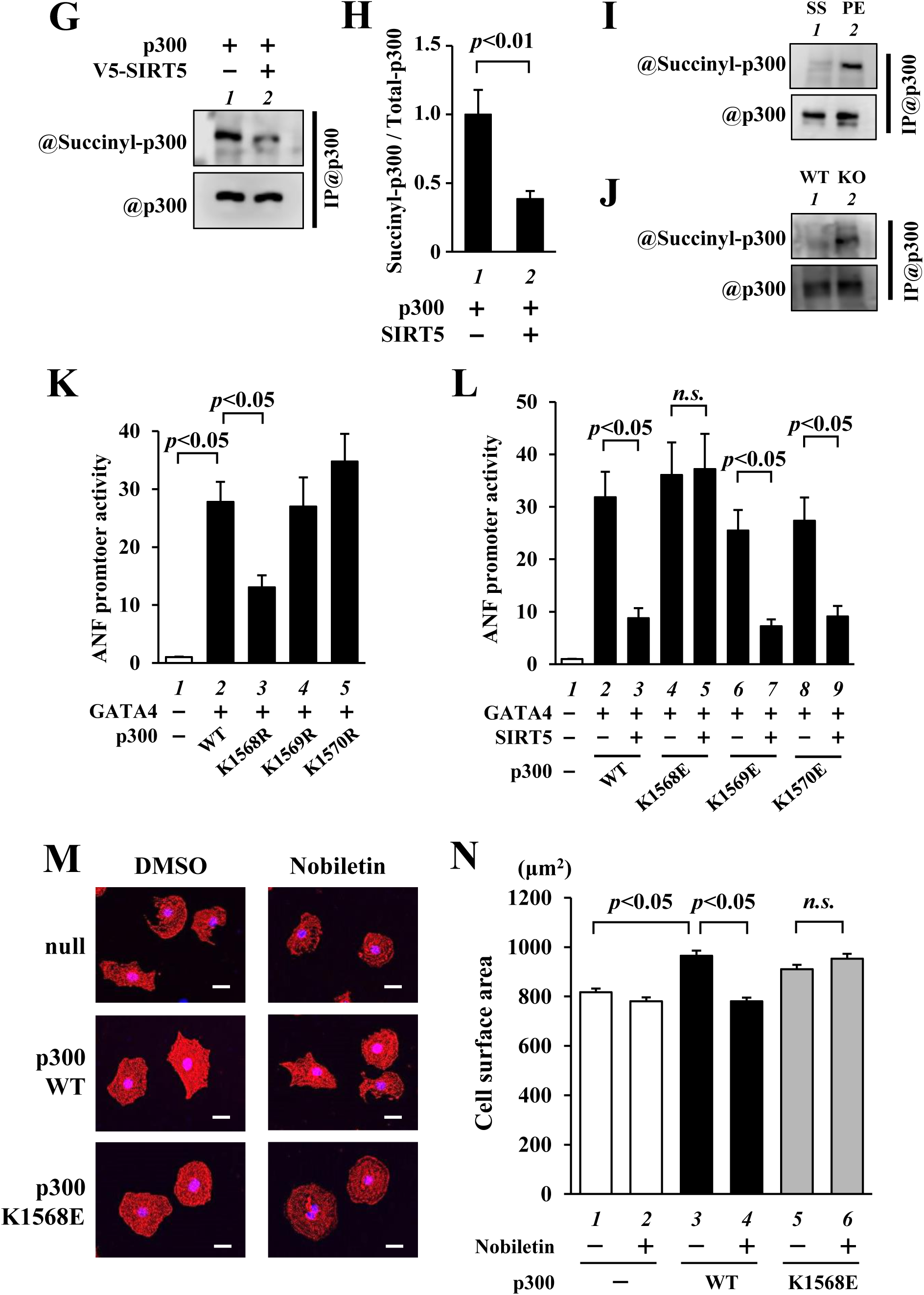
SIRT5 de-succinylates p300 and down-regulates p300-induced hypertrophic responses. **(A)** HEK293T cells were transfected with indicated vectors for 48 h. Nuclear extracts from these cells were subjected to immunoprecipitation using anti-FLAG M2 agarose, followed by western blotting. **(B)** The acetylated level of GATA4 was normalized to the total GATA4. **(C)** The binding level of p300 to GATA4 was normalized to precipitated GATA4. **(D)** HEK293T cells were transfected with pANF-luc and pRL-SV40 in the presence or absence of indicated constructs. The relative promoter activities were shown as means ± SEM from three independent experiments, each carried out in duplicate. **(E)** HEK293T cells were transfected with indicated constructs. Immunoprecipitated samples with anti-p300 or anti-V5 antibodies were subjected to western blotting. **(F)** The scheme of p300 functional domains and deletion mutants (upper panel). The interaction between V5-SIRT5 and various deletion mutants of p300 were shown on the right. Recombinant V5-SIRT5 prepared from *E.coli* BL21(CE3), was incubated with GST-fused deletion mutants of p300 for 2 h. Binding proteins were detected by western blotting (bottom panel). +: (binding), and −:(none). TAZ: transcriptional adaptor zinc-binding domain, KIX: kinase inducible domain interaction domain, Br: bromodomain, RING: RING finger domain, PHD: plant homeodomain, HAT: histone acetyltransferase, ZZ: ZZ type zinc finger domain, IBiD: Interferon-binding domain, CH: cysteine/histidine-rich domain. **(G)** HEK293T cells transfected with indicated constructs were treated with 1 µM TSA and 10 mM NAM for 6 h. Immunoprecipitated samples with anti-p300 antibody were subjected to western blotting. **(H)** The amount of succinyl-p300/total-p300 was calculated. The data are shown as means ± SEM from 3 independent experiments. **(I and J)** Nuclear extracts from cardiomyocytes stimulated with PE or saline **(I)** and hearts of SIRT5-KO and WT mice **(J)** were immunoprecipitated with anti-p300 antibody, followed by western blot using anti-succinyl lysine antibody. **(K and L)** HEK293T cells were transfected with pANF-luc, pRL-SV40, and indicated constructs. The relative promoter activities were shown as means ± SEM from three independent experiments, each carried out in duplicate. **(M)** Cultured cardiomyocytes were transfected with p300WT, p300K1568E, or null as control and incubated for 48 h. These cells were stained with an anti-β-MHC (red) antibody, and nuclei were labeled with Hoechst33258 (blue). Scar bar: 20 μm. **(N)** The surface area of β-MHC-positive cardiomyocytes was automatically measured. The data were shown as means ± SEM from three independent experiments.

Next, to examine whether SIRT5 binds to p300, HEK293T cells were transfected with p300 and V5-SIRT5, and immunoprecipitation western blotting was performed. SIRT5 interacted with p300 (Figure 7E). A GST pull-down assay showed that V5-SIRT5 was physically bound to p300 from aa1069 to 1459 (Figure 7F and Supplemental Figure S11B). This domain contains the RING finger and PHD domains, which negatively regulate p300-HAT activity by blocking the HAT active site^38,39^. As nobiletin enhances SIRT5-mediated desuccinylase activity, desuccinylation of p300 by SIRT5 may regulate p300-HAT activity and p300-induced hypertrophic responses. To determine whether p300 was succinylated, HEK293T cells expressing p300 in the presence or absence of SIRT5 were treated with 3 µM trichostatin A (TSA) and 50 mM nicotinamide (NAM) for 24 hours. Nuclear extracts were immunoprecipitated with an anti-p300 antibody, followed by western blotting with an anti-succinyl-lysine antibody. The results show that SIRT5 decreased p300 succinylation (Figures 7G and 7H). To investigate whether hypertrophic stimulation affects the p300 succinylation level, PE-stimulated cardiomyocytes were examined. PE stimulation increased the p300 succinylation levels in cultured cardiomyocytes (Figure 7I). Furthermore, cardiac succinylated p300 accumulated in the heart of SIRT5-KO mice compared to that of WT mice (Figure 7J).

To identify the p300 succinylation site, the p300 precipitate from HEK293T cells was subjected to mass spectrometric analysis^33^. LC-MS/MS spectrum analysis identified the sequence, including a p300 peptide (1566-1583), and one of the three lysines (1568-1570) was succinylated (Supplemental Figures S12A-S12B). To determine which succinylated lysine of p300 regulates p300-HAT activity, we mutated lysine to arginine (deficient in succinylation) or glutamic acid (a mimic of succinylation). A reporter gene assay demonstrated that only the succinylation-defective p300 K1568R mutant decreased p300-induced GATA4-dependent ANF and ET-1 promoter activation (Figure 7K and Supplemental Figures S13A and S13C). Moreover, the succinylation mimic mutant of p300 K1568E did not exhibit SIRT5-mediated inhibition of ANF or ET1 promoter activation (Figure 7L and Supplemental Figures S13B and S13D). To investigate whether p300 K1458 succinylation was involved in the inhibitory effect of nobiletin on cardiomyocyte hypertrophy, cultured cardiomyocytes were transfected with constructs expressing the p300 K1568E mutant and then treated with nobiletin. Immunofluorescence demonstrated that, although nobiletin treatment suppressed intact p300-induced cardiomyocyte hypertrophy, the p300 K1568E succinylation mimic mutant did not exhibit this inhibitory effect (Figures 7M and 7N). Taken together, these results suggest that SIRT5 desuccinylates p300 at the 1568 lysine residue and reduces p300-HAT activity. The p300 lysine residue is required for the hypertrophic response.

### Nobiletin/SIRT5 axis is involved in p300-induced histone H3K9 acetylation in cardiomyocytes

To investigate whether the nobiletin/SIRT5 axis regulates PE-induced acetylation of histone H3K9, a target of p300, cardiomyocytes were transfected with siSIRT5 and treated with nobiletin in the presence or absence of PE. As shown in Figure 8A, nobiletin treatment significantly suppressed the PE-induced acetylation of histone H3K9. However, SIRT5 knockdown disrupted this effect (Figure 8B). Conversely, SIRT5 overexpression prevented PE-induced histone H3K9 acetylation (Figures 8C and 8D). Furthermore, the overexpression of p300WT, but not of the p300 K1468E mutant, increased histone H3K9 acetylation (Figures 8E and 8F). These findings support the view that the nobiletin/SIRT5 axis exhibits antihypertrophic effects by downregulating p300-mediated HAT activity.

**Figure. 8.**
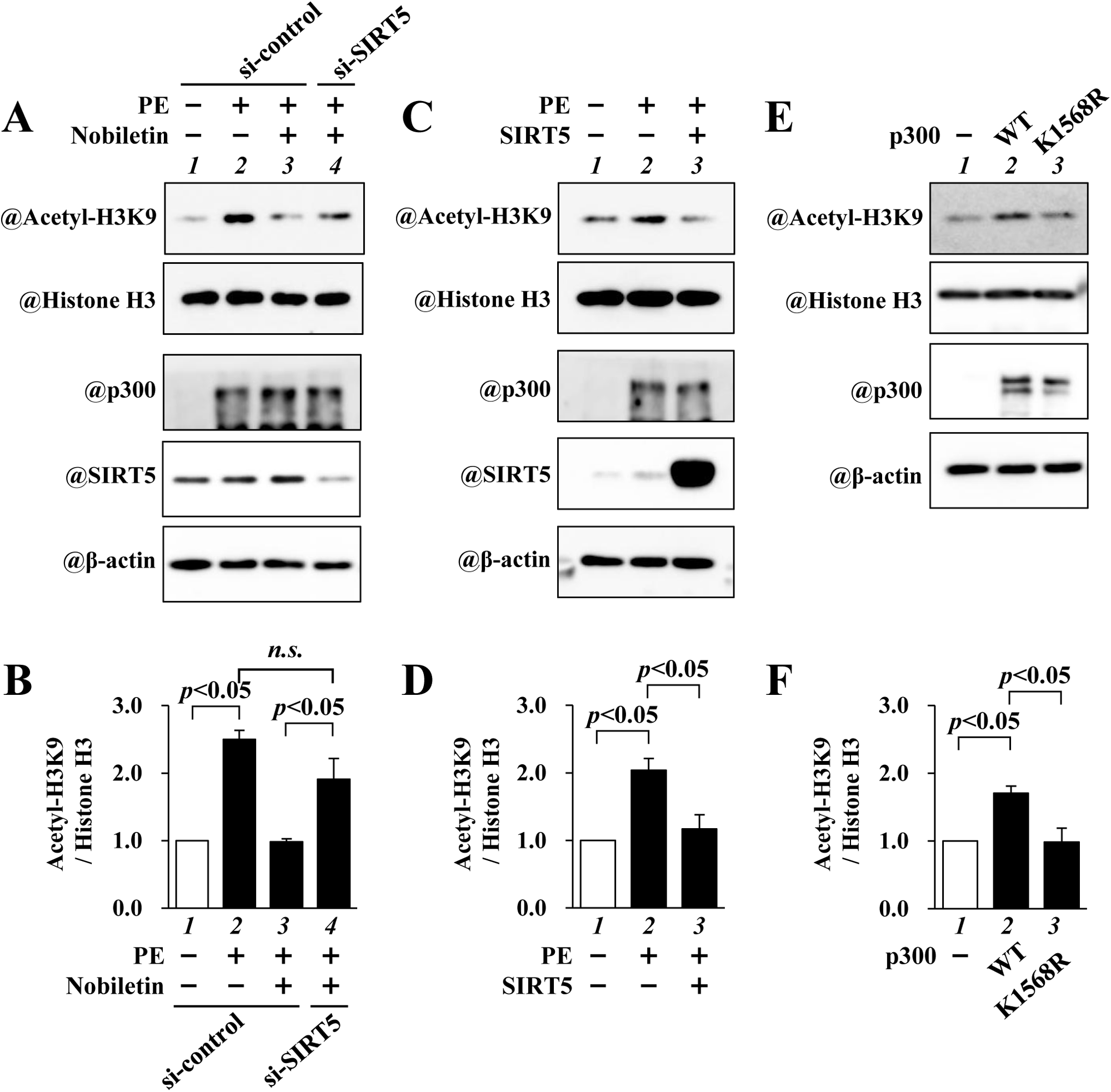
Nobiletin/SIRT5 axis is involved in the acetylation of Histone H3K9 in cardiomyocytes. **(A)** Cardiomyocytes were transfected with si-SIRT5 or si-control for 48 h, treated with nobiletin for 2 h, and stimulated with PE for 48 h. **(C)** Cardiomyocytes were infected with adenovirus encoding SIRT5 or LacZ as a control for 24 h and then stimulated with PE for 48 h. **(E)** Cardiomyocytes were transfected with p300WT, p300 K1568R, or null as a control for 2 h and cultured for 48 h. Nuclear extracts and acid extracts from these cells were subjected to western blotting. **(B, D, F)**. The amounts of acetylated H3K9/Histone H3 were calculated. The data are shown as means ± SEM from 3 independent experiments.

## Discussion

The current study demonstrated that nobiletin, a polymethoxyflavonoid, inhibits cardiomyocyte hypertrophy and exerts cardioprotective effects in two different animal models of heart failure. Proteomics identified SIRT5 as a novel target of nobiletin. We also demonstrated that SIRT5 is required for the inhibition of cardiac hypertrophy and systolic dysfunction induced by nobiletin. Furthermore, SIRT5 desuccinylated p300 and suppressed p300-HAT activity, thereby preventing hypertrophic responses and heart failure development. Our findings suggest that the nobiletin/SIRT5/p300 axis is a novel functional mechanism and potential therapeutic target for the treatment of heart failure.

A notable finding of this study is the identification of a novel target of nobiletin, SIRT5, which ameliorates cardiac hypertrophy and heart failure development. SIRT5 belongs to the sirtuin protein family and exhibits NAD-dependent lysine deacetylase activity^40^. Recently, SIRT5, a lysine desuccinylase and demalonylase^34^, was reported to play an important role in cardiovascular diseases^27–30^. SIRT5 regulates cardiac oxidative metabolism under pressure overload and improves survival^28^. SIRT5 protects against acute cardiac ischemia via a hepatic/cardiac crosstalk mechanism^27^. Furthermore, SIRT5 knockout mice have demonstrated ameliorated TAC-induced cardiac dysfunction^30^. These findings regarding the cardioprotective effects of SIRT5 contradict our findings. We analyzed systemic SIRT5-KO mice and found no differences in cardiac function or hemodynamics under physiological conditions between SIRT5-KO and WT mice (Supplemental Table S6). SIRT5-KO and WT mice exhibited no significant differences in TAC-induced cardiac dysfunction or hypertrophy (Figures 6B-6D). In both the MI and TAC models, SIRT5 mRNA levels decreased and the nuclear succinylation level increased in the heart (Supplemental Figure S7). It is possible that the effect of SIRT5-KO on cardiac hypertrophy in response to TAC was not observed because cardiac SIRT5 levels are diminished in chronic heart failure. Moreover, SIRT5 overexpression inhibits TAC-induced cardiac fibrosis^29^, and we confirmed a similar effect in SIRT5-TG mice with different genetic backgrounds. Nobiletin activates SIRT5 desuccinylation activity, induces p300 desuccinylation, and suppresses HAT activity, providing a novel mechanism to suppress cardiac hypertrophy and heart failure development. This mechanism is important for understanding the physiological activity of nobiletin.

Small-molecule compounds such as nobiletin exert their biological activity by binding to and modulating the activity of target molecules^24,25^. However, little is known regarding the target molecules of nobiletin. Target-fishing methods using biotin probes can detect a variety of proteins^41^. Based on MASS analysis, several candidate nobiletin target molecules were identified. In addition to SIRT5, the existing nobiletin target molecules *mkap1*, an extracellular signal-regulated kinase (ERK), and *prkaa1*, an AMP-activated protein kinase 1 (AMPK), were included (Supplemental Data 1). Therefore, our MASS results indicate reasonable candidate nobiletin target molecules. Nobiletin exerts anti-leukemic effects in K562 cells by activating ERK^42^. Nobiletin and its metabolites activate ERK and restore NMDA receptor antagonist-induced memory impairment^43^. Therefore, nobiletin may directly activate ERK. However, activation of the ERK pathway causes physiological hypertrophy^44^. This effect was inconsistent with the antihypertrophic effects of nobiletin observed in our study. AMPK acts as an intracellular energy sensor and has multifaceted cardioprotective effects by correcting low-energy states and improving the energy supply in failing hearts^45^. AMPK also negatively regulates the myocardial hypertrophic response via SIRT1 activation^46^. Nobiletin activates the AMPK signaling pathway and exerts antidiabetic effects^47^. Nevertheless, treatment with dorsomorphin, an AMPK inhibitor, and Ex527, a SIRT1 inhibitor, had no effect on the inhibitory effects of nobiletin on cardiomyocytes (Supplemental Figure S14). Based on these findings, we conclude that SIRT5, but not ERK or AMPK/SIRT1, plays a role in nobiletin-mediated antihypertrophic effects.

SIRT5 is highly expressed in the heart, liver, and brain^48^. SIRT5 is involved in the urea cycle and energy metabolism in the liver via mitochondrial deacetylation/desuccinylation^35,49^. SIRT5 ameliorates the progression of Alzheimer’s disease by activating autophagy^50^. However, few reports on the direct targets of SIRT5 exist. Although SIRT5 is mainly localized in the mitochondria and plays an important role in regulating lipid metabolism and mitochondrial pathways, some SIRT5 has been identified in the nucleus^51^. The present study also confirmed that SIRT5 was mainly localized in the nuclei of cardiomyocytes (Figure 4C). The level of nuclear succinylation increased in chronic heart failure with SIRT5 downregulation (Supplemental Figure S8), and SIRT5 regulated p300-HAT via p300 desuccinylation (Figure 7), suggesting that SIRT5 may be an important transcriptional regulator via nuclear succinylation in cardiomyocytes. Although SIRT5 does not contain a nuclear signal sequence^52^, several nuclear transport proteins such as Kpna3 and Kpna4, members of the importin family, were identified in the MASS analysis of nobiletin-binding proteins. Therefore, SIRT5 may be translocated to the nucleus via these proteins.

The ANF and ET-1 promoter constructs used in this study contained GATA-binding sites, and the transcription factor GATA4^57^ regulated their transcriptional activities. The present study showed that SIRT5 prevented p300/GATA4- and PE-induced promoter activation (Figures 1D and 7D, Supplemental Figures S1 and S12A), p300-mediated acetylation of GATA4 without altering the formation of the p300/GATA4 complex (Figures 7A–7 B), and PE-induced acetylation of histone H3K9 (Figures 8C-8D), suggesting that SIRT5 suppresses hypertrophic responses via the regulation of p300-HAT activity. In addition, the deacetylase activity of SIRT5 is weaker than its desuccinylase activity^53,54^. Our study demonstrated that nobiletin enhanced the desuccinylase activity of SIRT5, whereas its deacetylase activity was decreased (Figures 4B and Supplemental Figure S6C). Furthermore, SIRT5 decreased the succinylation level of p300 and p300-mediated GATA4 acetylation (Figures 7A-7B, 7G-7H). We also found that SIRT5 directly binds to the central region of p300 (amino acids 1069-1459)^39,55^ containing the bromodomain/PHD/RING module, which plays an important role in the regulation of p300-HAT activity and is a part of the HAT active module. This module negatively regulates p300-HAT activity by interacting with the lysine-rich autoinhibitory loop (AIL) domain^39,56^. These findings indicate that SIRT5 regulates p300-HAT activity through p300 desuccinylation.

Moreover, we identified the lysine residue at position 1568 as a novel p300 posttranslational modification site (Supplemental Figure S13 and 7) and found that SIRT5 could desuccinylate p300 at this residue (Figure 7G). This site is located at the base of the AIL domain, and the neighboring lysine residues are auto-acetylated by p300^39,56^. Although this site is not included in the p300 SIRT5-binding region (1069-1459), these sites are not far apart, given the crystal structure of p300^39^. Thus, it is reasonable to assume that SIRT5 can desuccinylate this site. The bromodomain/PHD/RING module interacts with the AIL domain and covers the substrate-binding site of p300-HAT^55^. This suggests that SIRT5 regulates p300-HAT activity by controlling the opening and closing of the cover via desuccinylation of the base of the AIL domain. PE-induced histone H3K9 acetylation was suppressed by nobiletin treatment and SIRT5 overexpression, and SIRT5 knockdown diminished the nobiletin-mediated inhibition of H3K9 acetylation (Figures 8A and 8C), supporting the hypothesis that the nobiletin/SIRT5 axis could regulate p300-HAT activity and hypertrophic responses. In contrast to deacetylation, desuccinylation is catalyzed by only SIRT5, and PE stimulation increases the level of p300 succinylation in cardiomyocytes (Figure 7I), and SIRT5 expression is decreased and nuclear succinylation levels are increased in hearts with chronic heart failure *in vivo* (Supplemental Figure S8), suggesting that the accumulation of p300 succinylation might further aggravate p300-HAT activity and heart failure development. Therefore, further analysis of the correlation between the heart failure stage and p300 succinylation levels is necessary. In the present study, we did not identify an enzyme capable of succinylating p300. Although Wagner et al. reported that protein succinylation occurs in an enzyme-independent manner^57^, Zorro et al. showed that p300 can succinylate H3K122 and destabilize the nucleosome structure^58^, indicating that p300 may regulate its HAT-activity by self-succinylation. This study suggests that the nobiletin/SIRT5 axis is a unique pathway that intervenes in p300 succinylation.

## Limitations

In this study, we could not clarify how nobiletin enhances the enzymatic activity of SIRT5. Resveratrol, a SIRT1 activator, binds to the active pocket of SIRT1 and enhances enzymatic activity^59^. Therefore, a similar mechanism is expected for nobiletin/SIRT5. Commercially available anti-p300 antibodies have very poor immunoprecipitation efficiency for p300 in tissue samples, and we were unable to detect changes in p300 succinylation following TAC or MI. Overcoming this problem and proceeding with further p300 analysis is critical.

## Conclusion

The present study demonstrated that the polymethoxyflavonoid nobiletin prevents cardiomyocyte hypertrophy *in vitro* and TAC- and MI-induced myocardial hypertrophy and cardiac dysfunction *in vivo*. We also found that the SIRT5/p300 axis is involved in the therapeutic potency of nobiletin. This novel regulatory mechanism of nobiletin is valuable for understanding its beneficial physiological activity. This study provides novel strategies for the treatment of heart failure using nobiletin.

## Author contribution

Sunagawa Y: Conceptualization, Methodology, Investigation, Writing - Original Draft & Editing; Funamoto M and Toshihide Hamabe-Horiike, Investigation and Data Curation; Hieda K, Yabuki S, Tomino M, Ikai Y, Suzuki A, Ogawahara S, Yabuta A, Sasaki H, Ebe A, Naito S, and Takai H: Investigation, Visualization, Formal analysis; Shimizu K, Shimizu S, Kawase Y, and Naruta R: Investigation; Katanasaka Y: Data Curation; Asakawa T, Kan T, Mori K, Murakami A, Ogura M, Inagaki J and Koji Hasegawa: Resources, Supervision; Morimoto T: Project administration, Writing - Original Draft & Editing.

## Acknowledgments

We thank Sayuri Miyagishima, Hiroko Mochizuki, and Nobuko Okamura for their technical assistance. We thank Editage (www.editage.com) for English language editing.

## Ethical statement

This study approved by to the Guide for the Care and Use of Laboratory Animals by the Institute of Laboratory Animals, Kyoto Medical Center (KMC2-25-4) and University of Shizuoka (US176261) in Japan.

