## Supplemental Materials and Methods for "Nobiletin, a Polymethoxyflavonoid, Activates the Desuccinylase Activity of SIRT5 and Prevents the Development of Heart Failure"

### ***Materials***

Purified Nobiletin (98%>) was obtained from Prof. Akira Murakami (University of Hyogo, Japan)[1]. Phenylephrine (PE); an α1 agonist, Trichostatin A (TSA); a pan-histone de-acetylase inhibitor, nicotinamide (NAM); a Sirtuin inhibitor, dorsomorphin; an AMP-activated protein kinase (AMPK) inhibitor, EX527; a SIRT1 inhibitor, and dimethyl sulfoxide (DMSO) were purchased from FUJIFILM Wako Pure Chemical, Osaka, Japan. Nobiletin, dorsomorphin, and EX527 were dissolved in DMSO. PE and NAM were dissolved in distilled water. TSA was dissolved in 99.5% ethanol. These compounds were stored in a -20ºC freezer. For animal experiments, Nobiletin was dissolved in 1% gum arabic solution before treatment. Biotin-conjugated Nobiletin (bio-Nobi) and TokyoGreen-conjugated Nobiletin (TG-Nobi) were synthesized by Prof. Toshiyuki Kan and Tomohiro Asakawa (University of Shizuoka, Japan)[2].

### ***Animal experiments***

Sprague-Dawley (SD) rats and C57BL6J mice were purchased from Japan SLC Inc, Shizuoka, Japan. SIRT5-TG mice were gifted from Prof. Nobuya Inagaki (Kyoto University, Japan)[3]. SIRT5-KO (129 background) and control wild-type (WT) mice were obtained from Jackson Laboratory (Bar Harbor, ME, stock#012757) and backcrossed to C57BL/6. All animal experiments conformed to the Guide for the Care and Use of Laboratory Animals by the Institute of Laboratory Animals, Kyoto Medical Center (KMC2-25-4) and University of Shizuoka (US176261). All mice and rats used in our experiments were maintained under 12 h light/dark lighting conditions (lights on at 8:00 A.M.) with food and water available *ad libitum*. For genotyping, genomic DNA was extracted from the mouse tail. The tails were incubated in 180 μL lysis solution (50 mM NaOH, 1 mM EDTA) for 10 min at 95ºC, followed by the addition of 20 μL 1 M Tris-HCl pH8.0. Their supernatants were used as a template for genotyping PCR. All transgenic mice in our experiments were genotyped with specific primers. The genotyping primers are shown in Supplemental Table S1.

The mice at 8 weeks old were anesthetized with avertin and then subjected to TAC surgery. The surgery was performed as described in the previous study[4]. A sham surgery was performed with the same protocol, except the suture around the aortic arch was not ligated. One day after surgery, the mice were randomly divided into two groups; vehicle as a control (1% gum arabic) and Nobiletin (20 mg/kg/day), and orally administrated by gastric gavage for 8 weeks. Myocardial infarction (MI) was generated in SD rats at eight weeks old by ligating the proximal left anterior descending (LAD) coronary artery through a left thoracotomy, as described previously[5]. Only a left thoracotomy was performed in sham-operated rats. At one week after LAD ligation, moderated MI rats (FS<40%) were randomly assigned to 3 groups; vehicle as a control, the low-dose (2 mg/kg/day), and the high-dose of Nobiletin (20 mg/kg/day). Oral administrations were repeated for 7 weeks. [65]

***Analysis of Echocardiographic and Hemodynamic Parameters***

The cardiac function of all rats and mice were noninvasively evaluated by echocardiography according to the methods described previously[7]. In brief, images were recorded using a 10- to 12-MHz phased-array transducer (model 21380A with HP SONOS 5500 imaging system, Philips, Amsterdam, Nederland). LV internal dimensions of diastole and systole (LVIDd and LVIDs), LV posterior wall thickness (PWT), fractional shortening (FS), and heart rate (HR) were measured with M-mode tracings from the short-axis view of the LV at the papillary muscle level. All measurements were performed in a blinded fashion according to the guidelines of the American Society for Echocardiology and averaged over three consecutive cardiac cycles. Systolic blood pressure (SBP) and diastolic blood pressure (DBP), and pulse rate (PR) were measured in all rats and mice by the tail-cuff method according to the manufacturer's instructions (BP98-A, Softron. Inc, Tokyo, Japan).

***Histological analysis***

The excised hearts were cut into two transverse slices at the mid-level of papillary muscles, fixed in 3.7% paraformaldehyde (PFA) and embedded in paraffin. Heart tissues were sliced into 4-μm-sections and stained with hematoxylin and eosin (HE) and Masson trichrome (MT). Quantitative assessments of cross-sectional myocardial cell diameter and perivascular fibrosis area were described previously[8].

***Quantified Polymerase Chain Reaction (QPCR)***

Total RNA from heart tissue and cardiomyocytes was extracted using Trizole Reagent (Thermo Fisher Scientific, MA, USA). cDNA was synthesized from 1 μg of total RNA using ReverTra Ace qPCR RT Master Mix (TOYOBO, Osaka, Japan). The sequences of used primers were described in Supplemental Table S1. The amplification of the gene was quantified by LightCycler 96 Real-Time PCR System (Roche, Basel, Switzerland) with KOD SYBR qPCR Mix (TOYOBO, Osaka, Japan). The protocols were performed according to the manufacturer's instructions. 18S rRNA was used as a reference gene, and relative activity was calculated by the ΔΔCT method. The data in the control group are set to a relative value of one, and each value is represented as the mean ± standard error.

***Plasmid constructs***

Human SIRT5 cDNA with V5 tag was inserted into pENTR-1A and cloned into pDEST17, pcDNA3.2-DEST, and pAd-CMV-DEST by the GATEWAY system (Thermo Fisher Scientific, MA, USA). pcDNA3.2-null and pAd-CMV-GFP were used as a controls. The fragments of Human p300 (1-450, 419-761, 762-1098, 1069-1459, 1284-1673, 1514-1877, 1877-2160, 2160-2378) were inserted into pGEX-6p1. The expression vectors, pcDNA-null, pCMV-p300, pcDNA-FLAG-GATA4, pANF-luc, pET-luc, and pRL-SV40 were previously described[9]. According to the manufacturer's instructions, KOD-Plus-Mutagenesis (TOYOBO,

Osaka, Japan) was used for site-direct mutagenesis. All the cloned sequences were confirmed by DNA sequencing.

***Antibodies***

Commercial antibodies were used as follows: rabbit anti-SIRT5 (#8779), rabbit anti-acetyl-lysine (#9441), rabbit acetyl-Histone H3K9 (#9649), rabbit Histone H3 (#3399) antibodies (Cell Signaling Technology, MA, USA), rabbit anti-pan-succinyl lysine antibody (PTM-401, PTM Biolabs, IL, USA), goat and mouse anti-GATA4 (G-4, C-20), rabbit anti-V5 (C-9), anti-HA (F-7), and rabbit anti-p300 (N-15 and C-20) antibodies (Santa Cruz Biotechnology, TX, USA), and mouse anti-β-actin antibody (A5441, Sigma-Aldrich, MO, USA) were used as primary antibodies. Horseradish peroxidase-conjugated anti-mouse, anti-rabbit, and anti-goat antibodies (Santa Cruz Biotechnology, TX, USA) were used as secondary antibodies. Mouse anti-α-actinin antibody (Sigma-Aldrich, MO, USA), Alexa Fluor 555 conjugated goat anti-mouse IgG antibody, and Dylight649-conjugated goat anti-rabbit IgG antibodies (Thermo Fisher Scientific, MA, USA) were used for immune-staining.

***Cell culture, treatment, transfection, and infection***

HEK293T cells and HEK293A cells were maintained in Dullbecco's modified Eagle's medium (D-MEM) (Nakalai tesque, Kyoto, Japan) supplemented with 10% fetal bovine serum (FBS), penicillin, and streptomycin (Nakalai tesque, Kyoto, Japan). According to the manufacturer's instructions, these cells were transfected with expression plasmids using Polyethyleneimine MAX (Polysciences, PA, USA). For the production of recombinant adenovirus particles, the recombinant adenoviral DNAs encoding SIRT5 and GFP as a control were transformed into HEK293A cells. After seven days post-transfection, the recombinant adenoviruses were corrected and amplified in HEK293A cells. The adenovirus titer was determined by the median tissue culture infectious dose (TCID50) and estimated at $1.0 \times 10^9$

PFU/mL.

Primary neonatal rat cardiomyocytes were prepared as previously described[10]. In brief, hearts from 2-day-old SD rats were digested with 3 mg/mL collagenase type 2 (Worthington, NJ, USA) and 3 mg/mL pancreatin (Sigma-Aldrich, MO, USA) at 37°C in a water bath for 10 min. The supernatants containing cardiomyocytes were centrifugated at 1,500 g for 10 min and resuspended in D-MEM containing 10% FBS. After seven digestions, cells were plated on 10-cm culture dishes for one h to remove adherent non-cardiomyocytes. Non-adherent cardiomyocytes were plated at $2.5 \times 10^4$ cells/cm$^2$ on culture dishes or plates at 37°C in an incubator with 5% $CO_2$. After 24 h, cardiomyocytes were washed three times in D-MEM and used for experiments.

Cardiomyocytes were treated with or without Nobiletin for 2 h and then stimulated with 30 μM PE for 48 h. For a reporter assay, cardiomyocytes were transfected with the reporter constructs and expression plasmids using Lipofectamine LTX and Plus reagent (ThermoFisher Scientific, MA, USA) for 2 h and then stimulated with PE for 48 h. Firefly and sea pansy luciferase activities were measured by Dual-Luciferase® Reporter Assay (Promega, WI, USA), and the relative promoter activities were calculated as the ratio of firefly to sea pansy luciferase. Cardiomyocytes were infected with the recombinant adenovirus at an MOI of 10 for 24 h, and then the medium was changed to fresh serum-free media. After 48 h post-infection, cardiomyocytes were stimulated with PE for 48 h. For RNA interference, siRNA-control (si-control) and siRNA-SIRT5 (si-SIRT5) were purchased from Sigma-Aldrich (MO, USA). Cardiomyocytes were transfected with siRNA using Lipofectamine RNAi MAX according to the manufacturer's instructions (ThermoFisher Scientific, MA, USA). At six h after transfection, the medium was changed to fresh serum-free media. After 48 h post-transfection, cardiomyocytes were treated with or without Nobiletin for 2 h and subsequently stimulated with PE for 48 h.

***Immunofluorescent staining and measurement of cell surface area***

The cultured cardiomyocytes grown in 24 well plates (Thermo Fisher Scientific, MA, USA) were fixed in 3.7% PFA and stained using mouse anti-α-actinin antibody and Alexa555-conjugated anti-mouse antibody as previously described[11]. The α-actinin-positive cardiomyocytes were scanned with an ArrayScan™ High-Content Systems (Thermo Fisher Scientific, MA, USA), and the surface area of 300 cells was measured automatically using HCS Studio™ 2.0 Client Software (Thermo Fisher Scientific, MA, USA). The representative immunofluorescence staining images of cardiomyocytes were photographed using BZ-X810 (KEYENCE, Osaka, Japan).

***Purification and identification of Nobiletin binding proteins***

The preparation of cell lysates from heart tissue has been described previously[12]. Eighty mg lysate from rat hearts was preincubated with streptavidin sepharose (Thermo Fisher Scientific, MA, USA) for 1 h at 4°C. Then, the collected supernatant was incubated with biotinylated Nobiletin (bio-Nobi, 4 μM) or biotin (4 μM). After incubation for 1 h at 4°C, proteins associated with bio-Nobi or biotin were precipitated with streptavidin sepharose for 1 h at 4°C. After washing five times in 0.1 buffer (0.1 M KCl, 20 mM Trs-HCL pH 8.0, 1 mM $MgCl_2$, 10% Glycerol, and 0.1% Tween-20) containing 1 mM PMSF, 1 mg/ml pepstatin, 1 mM $Na_3VO_4$, 1 mM NaF, and 10 mM sodium butyrate, the binding proteins were eluted with 1 M NaCl, concentrated by StrataClean Resin (Agilent, USA), separated by NuPAGE-gel, and visualized by silver staining kit or Quick-CBB kit (FUJIFILM Wako Pure Chemical, Osaka, Japan). Mass spectrometry analysis was performed at the Taplin Biological Mass Spectrometry Facility, Department of Cell Biology, Harvard Medical School[13]. The Nobiletin binding protein was defined as MASS analysis data of bio-Nobi without those of biotin.

For *in vitro* binding assay, recombinant SIRT5 was expressed in *E.coli* BL21(DE3). Recombinant SIRT5 was added with or without 40 μM Nobiletin as a competitor and incubated in the presence or absence of 4 μM bio-Nobi for 2 h at 4°C. The binding proteins were precipitated with streptavidin sepharose, washed five times in 0.1 buffer, and subjected to western blotting. The band was photographed by LAS1000 plus (Cytiva, MA, USA).

***Western blotting and immunoprecipitation (IP)***

Nuclear extracts and acid extracts were described previously[9]. The total protein concentrations were measured using a Bradford Protein assay kit (BioRad, CA, USA). The equal amounts of proteins were separated on SDS-PAGE and transferred onto nitrocellulose membranes (Merck, Darmstadt, Germany) using a Mini Trans-Blot® cell (BioRad, CA, USA). The transfer was performed at a constant ampere setting of 200 mA for 2 h at 4°C. The membranes were blocked in 2% skim milk for 1 h at room temperature and reacted in PBS containing 0.1% Tween-20 (PBS-T) with primary antibodies at 1:1000 dilution for 2 h at room temperature. After washing three times for 10 min in PBS-T, the membranes were incubated in PBS-T with secondary antibodies at 1:2000 for 2 h at room temperature. The membranes were washed three times for 10 min in PBS-T and incubated in ImmunoStar LD (FUJIFILM Wako Pure Chemical, Osaka, Japan). The chemiluminescence from membranes was detected by LAS1000 plus and quantified by Multi Gauge V3.0 (Cytiva, MA, USA). For an IP of flag-tagged proteins, nuclear extracts were incubated with anti-flag M2 affinity agarose gel for 2 h at 4°C. For an IP of p300, nuclear extracts were incubated with 1 μg anti-HA, anti-p300, or anti-IgG antibodies for 2 h at 4°C and then added with 10 μL Protein A sepharose (Cytiva, MA, USA). After incubation for 2 h, precipitated samples were washed three times in 0.1 buffer, eluted by 0.1 M Glycine pH 2.5, and subjected to western blotting.

***In vitro de-succinylase and de-acetylase activities***

The de-succinylase and de-acetylase activities of SIRT5 *in vitro* was determined using the Fluorogenic SIRT5 and SIRT1 Assay Kits (BPS Bioscience, CA, USA), according to the manufacturer's instructions. In brief, human SIRT5 (300 ng) was pre-incubated with 100 μM Nobiletin, 100 μM Resveratrol, and DMSO as a control for 30 min, and then added reaction mixture for 15 min at room temperature. The de-succinylase or de-acetylase activities were measured by microplate reading fluorimeter at a wavelength in the range of 350-380 nm and detection of emitted light in the range of 440-460 nm. The data in the control group are set to a relative value of one, and each value is represented as the mean ± standard error.

***Identification of succinylated lysine of p300***

HEK293T cells were transfected with expression plasmids encoding p300 and SIRT5 and incubated for 24 h. Cells were treated with 1 μM TSA and 10 mM NAM for 24 hours to prevent de-acetylase and de-succinylase activity. Twenty mg nuclear extracts from these cells were immunoprecipitated with an anti-p300 antibody. Precipitated samples were eluted in 1xNuPAGE sample buffer, separated by NuPAGE-gel. Precipitated proteins were stained by a Quick-CBB kit (FUJIFILM Wako Pure Chemical, Osaka, Japan). The gel was cut and analyzed by Taplin Biological Mass Spectrometry Facility, Department of Cell Biology, Harvard Medical School[13].

***Statistical analysis***

Data are presented as the mean ± SEM. Statistical comparisons were performed using unpaired 2-tailed Student *t*-tests or ANOVA with Tukey-Kramer when appropriate, with a probability value $<0.05$ taken to indicate significance.
