## Supplemental figure for "Nobiletin, a Polymethoxyflavonoid, Activates the Desuccinylase Activity of SIRT5 and Prevents the Development of Heart Failure"

### Supplemental Figure S1

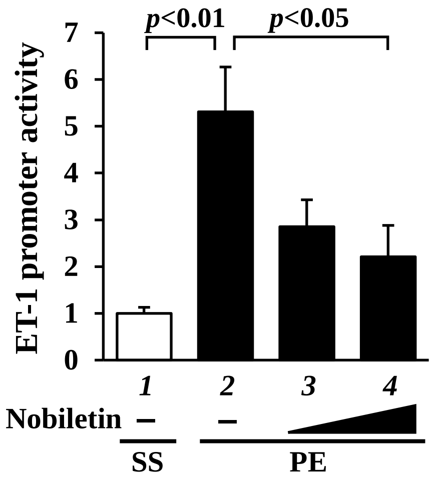


**Supplemental Figure. S1 Nobiletin significantly prevents PE-induced ET-1 promoter activation in cultured cardiomyocytes**

Cardiomyocytes were transfected with pET-luc and pRL-SV40 in the presence or absence of Nobiletin for 2 h, and then stimulated with PE for 48 h. The relative ET-1 promoter activity was calculated from the ratio of firely luciferase activity to sea pansy luciferase activity. The values were shown as means ± SEM from three independent experiments, each carried out in duplicate.

### Supplemental Figure S2

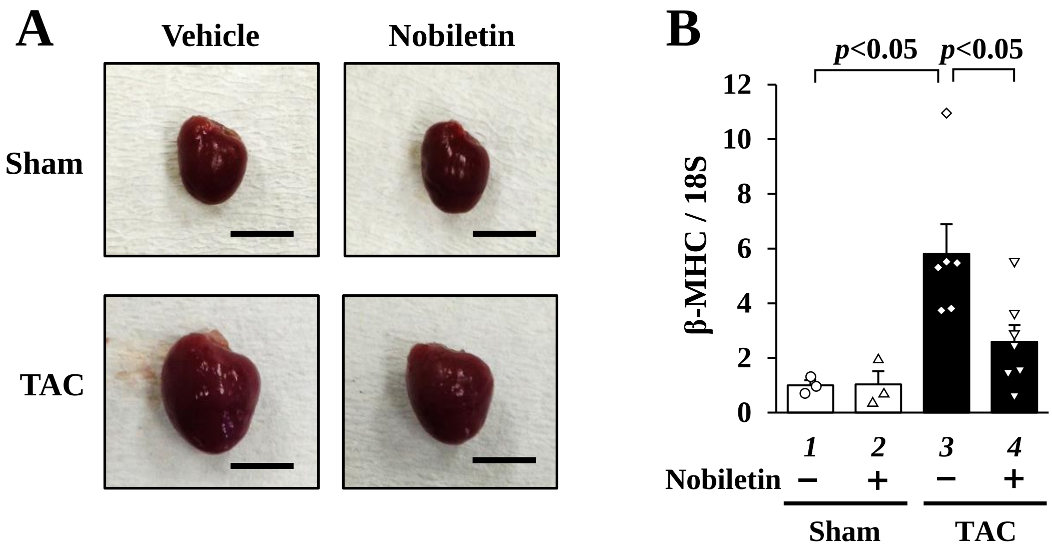


**Supplemental Figure. S2 Nobiletin significantly prevents pressure overload-induced cardiac hypertrophy in mice**

**(A)** The representative images of the whole heart from indicated groups at 8 weeks after surgery. Scale bar indicates 5 mm. **(B)** Total RNA from each heart was subjected to quantified RT-PCR. The mRNA level of β-MHC was normalized by that of 18S. The values were shown as mean ± SEM.

### Supplemental Figure S3

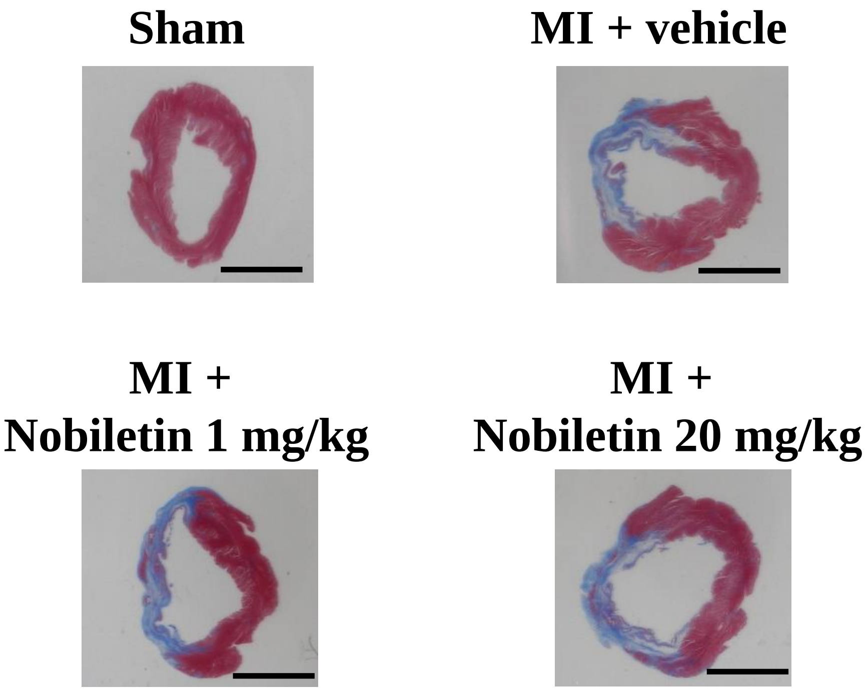


**Supplemental Figure. S3 The infarct size of the left ventricle was not changed by Nobiletin treatment in rats**

The representative images of MT-stained cross-sectional left ventricle from indicated groups at 8 weeks after surgery. The scale bar indicates 10 mm.

### Supplemental Figure S4

**A**

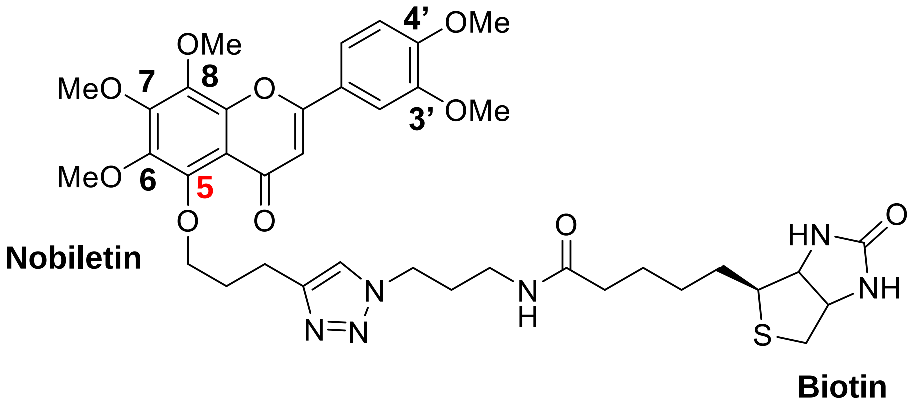


**B**

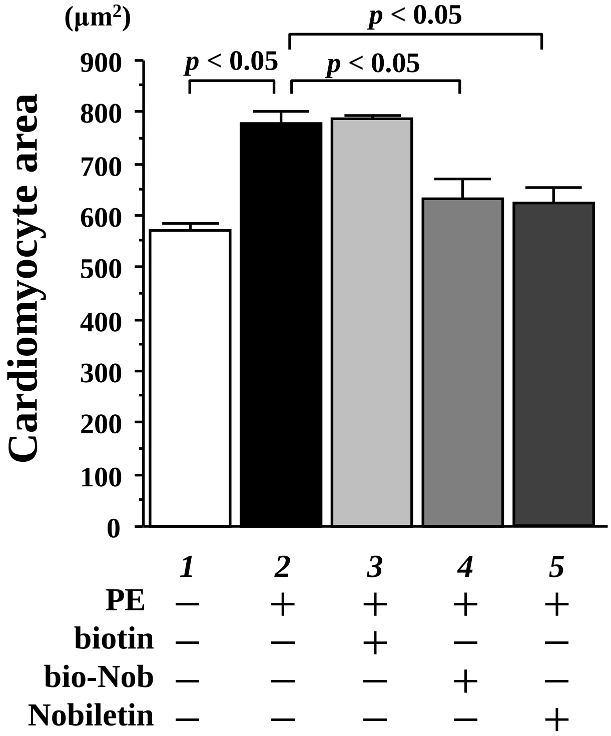


**Supplemental Figure. S4 Biotin-conjugated Nobiletin possesses an anti-hypertrophic effect**

**(A)** The structure of biotin-conjugated Nobiletin (bio-Nobi). **(B)** Cardiomyocytes were treated with Nobiletin, biotin, bio-Nobi, or DMSO as a control and stimulated with 30 μM PE for 48 h. The cell surface area of β-MHC-positive cardiomyocytes was measured. The data were shown as means ± SEM from three independent experiments.

### Supplemental Figure S5

A

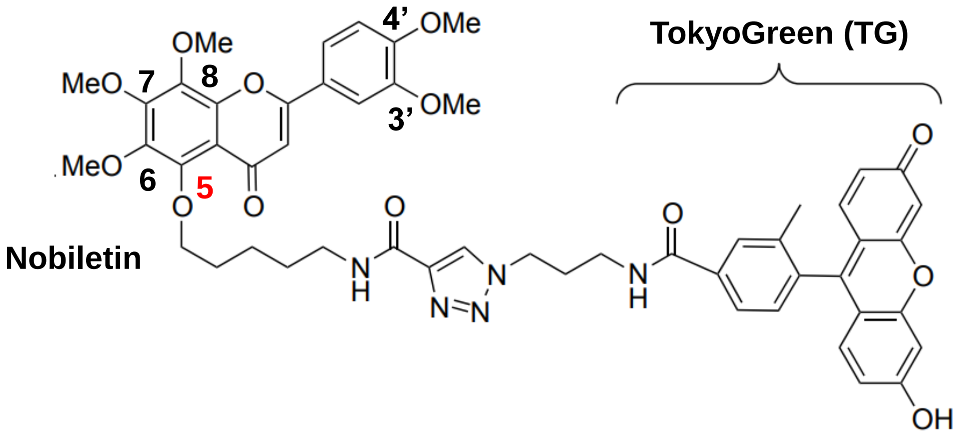


B

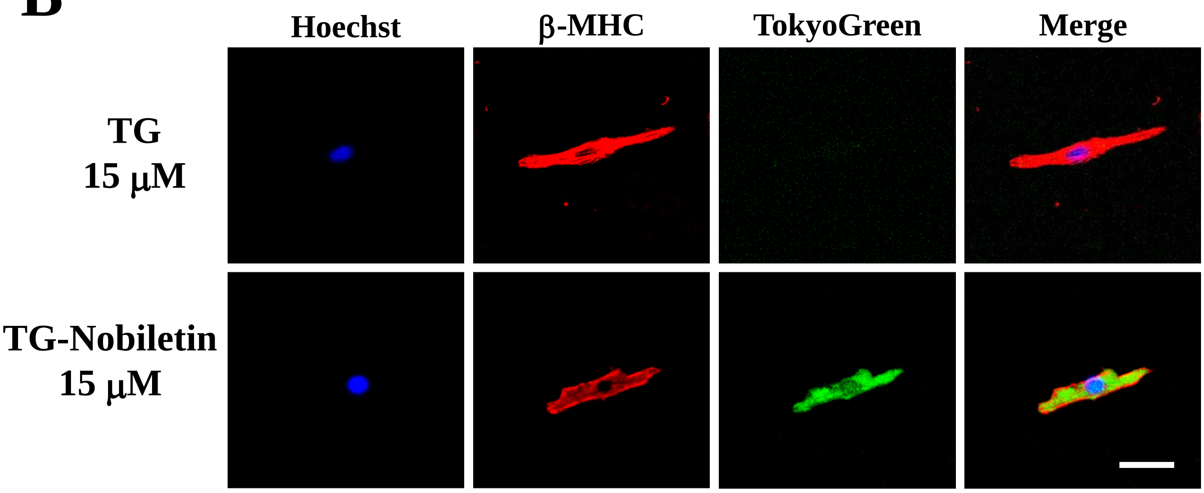


**Supplemental Figure. S5 Nobiletin was observed in the nucleus and cytosol of cardiomyocytes** **(A)** The structure of TokyoGreen-conjugated Nobiletin (TG-Nobi). **(B)** Cardiomyocytes were treated with TG or TG-Nobi for 2 h. These cells were stained with anti-β-MHC (red) antibody and Hoechst33258 (blue). Green signals indicated the distribution of Nobiletin.

### Supplemental Figure S6

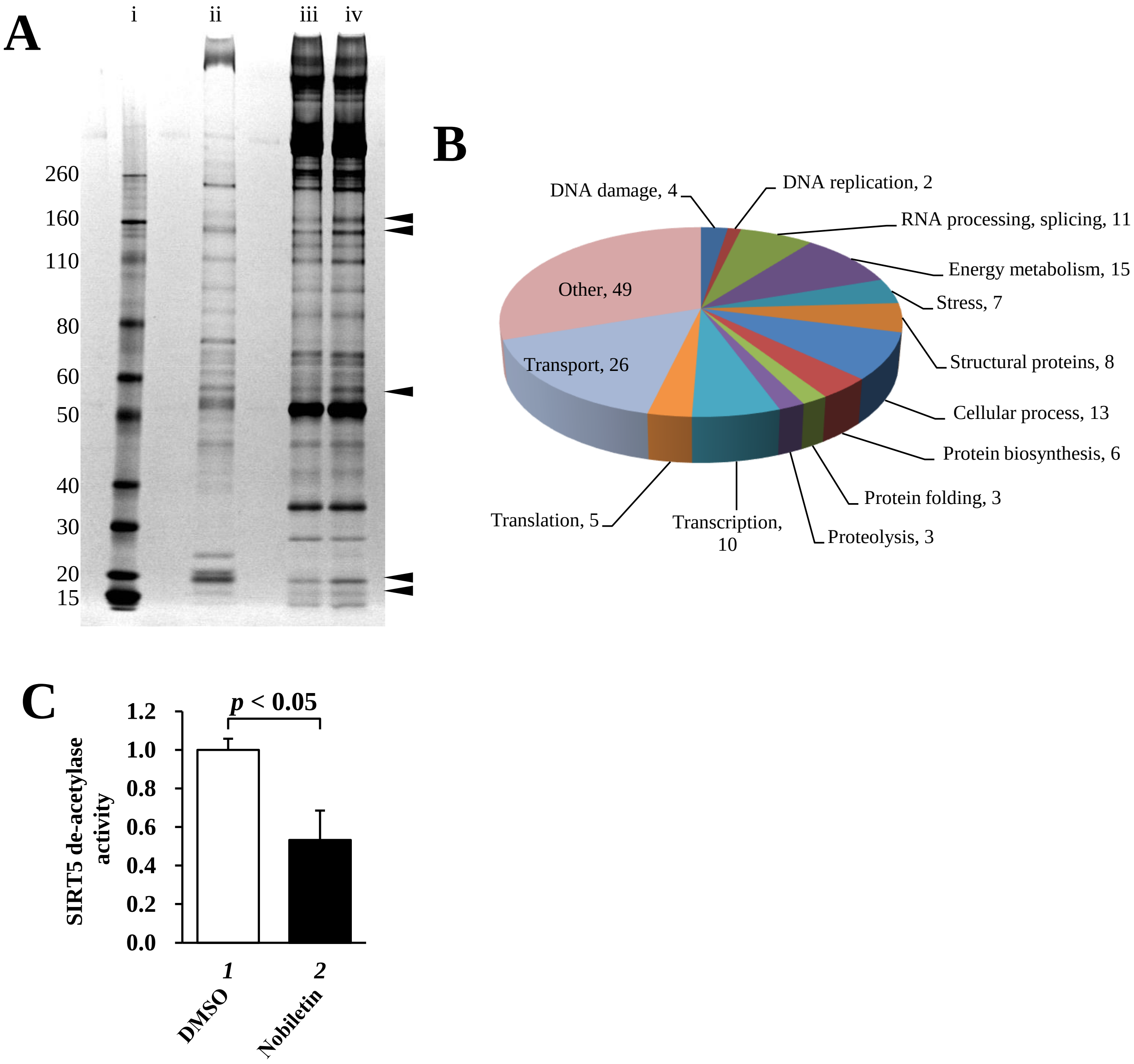


**Supplemental Figure. S6 Identification of Nobiletin binding proteins by a proteomic approach**

**(A)** Silver staining of Nobiletin binding proteins purified from rat heart. i: protein marker. ii: input. iii: biotin-binding proteins. iv: Nobiletin binding proteins. Detailed data were provided in Supplemental data1. **(B)** The classification of Nobiletin binding proteins. **(C)** The de-acetylation activities of SIRT5 in the presence or absence of Nobiletin (100 μM) were measured using a commercial kit. The relative values were shown as means ± SEM from three independent experiments.

### Supplemental Figure S7

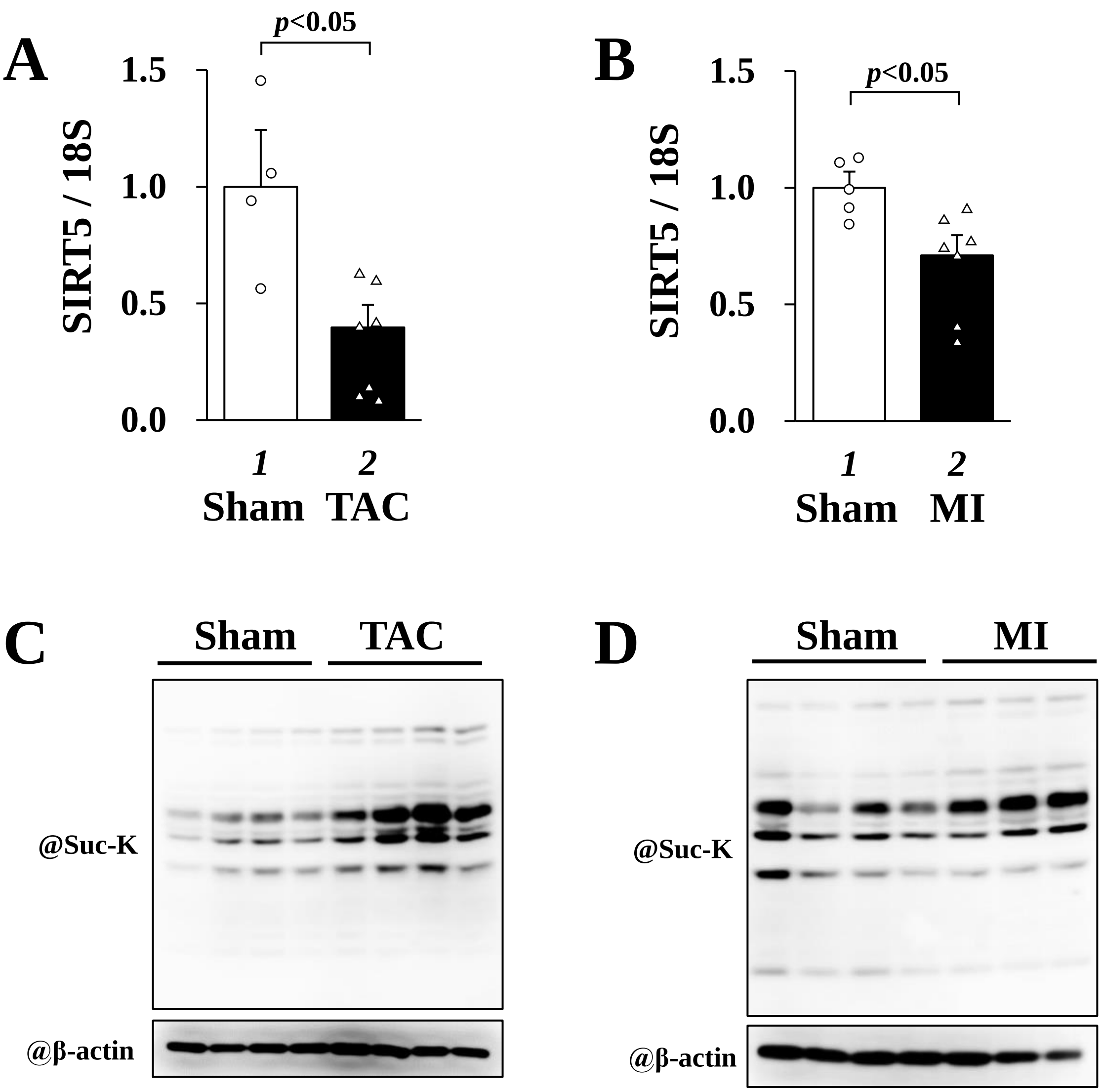


**Supplemental Figure. S7 Cardiac hypertrophic stimulations reduce mRNA levels of SIRT5 and accumulate the succinylated level of nucleus in the heart**

**(A and B)** Eight weeks-old C57BL6j mice were subjected to TAC or Sham surgery. Eight weeks-old SD rats were subjected to MI or Sham surgery. At 8 weeks after surgery, total RNAs from these hearts were subjected to quantified RT-PCR. The mRNA levels of SIRT5 were normalized by that of 18S. The data were shown as means ± SEM. **(C and D)** Nuclear extracts from these hearts were subjected to western blotting using an anti-Succinylated lysine antibody.

### Supplemental Figure S8

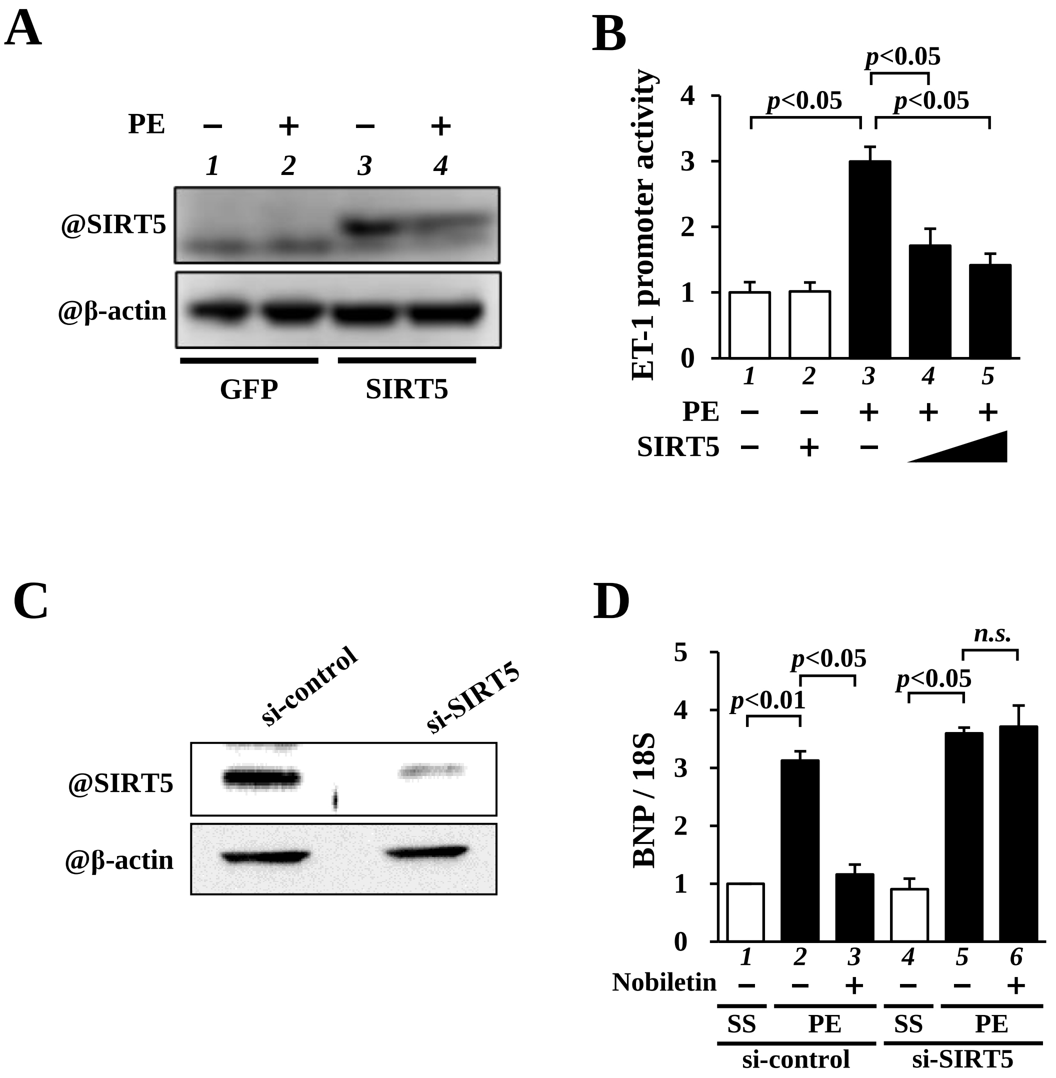


**Supplemental Figure. S8 The overexpression and knockdown of SIRT5 in cardiomyocytes were confirmed**

**(A)** Cardiomyocytes were infected with adenovirus encoding V5-tagged SIRT5 or LacZ as a control and stimulated with the presence or absence of PE for 48 h. Nuclear extract from these cells was subjected to western blotting. **(B)** Cardiomyocytes were transfected with pET-luc, pRL-SV40, and pcDNA-SIRT5 in the presence or absence of PE for 48 h. The relative promoter activities were shown as means ± SEM from three independent experiments, each carried out in duplicate. **(C)** Cardiomyocytes were transfected with si-SIRT5 or si-control as a control for 48 h. Nuclear extract from these cells was subjected to western blotting. **(D)** Cardiomyocytes were transfected with si-SIRT5 or si-control for 48 h, treated with Nobiletin for 2 h and then stimulated with PE for 48 h. Total RNAs from indicated cardiomyocytes were subjected to quantified RT-PCR. The mRNA level of ANF was normalized by that of 18S. The data were shown as means ± SEM from three independent experiments.

### Supplemental Figure S9

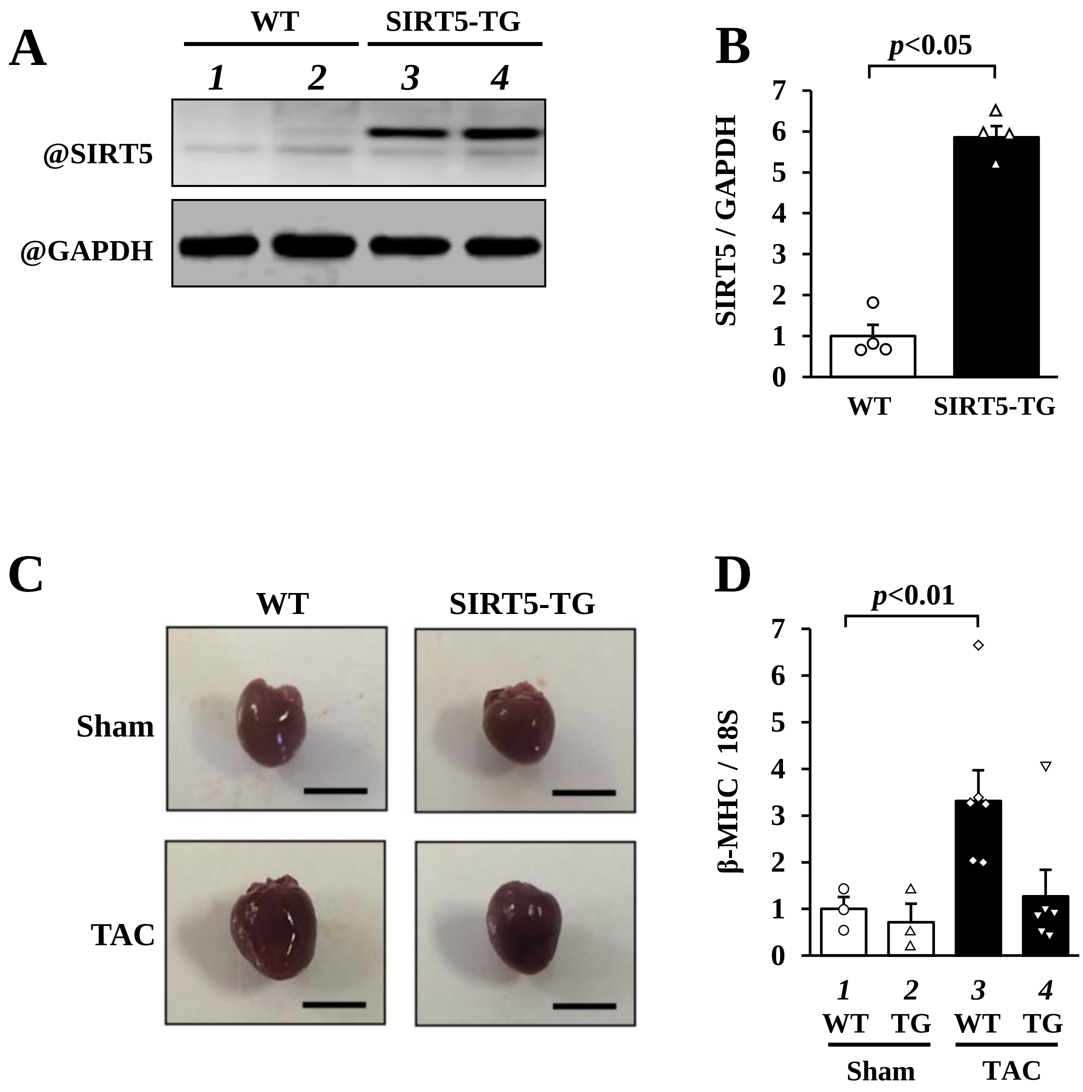


**Supplemental Figure. S9 Overexpression of SIRT5 prevents TAC-induced cardiac hypertrophy in mice**

**(A)** Nuclear extracts from the hearts of SIRT5-TG and WT mice were subjected to western blotting. **(B)** The expression level of SIRT5 was normalized to GAPDH. The data were shown as means ± SEM. **(C)** The representative images of the whole heart from indicated groups at 8 weeks after surgery. Scale bar indicates 10 mm. **(D)** Total RNA from each heart was subjected to quantified RT-PCR. The mRNA level of β-MHC was normalized by that of 18S. The values were shown as mean ± SEM.

### Supplemental Figure S10

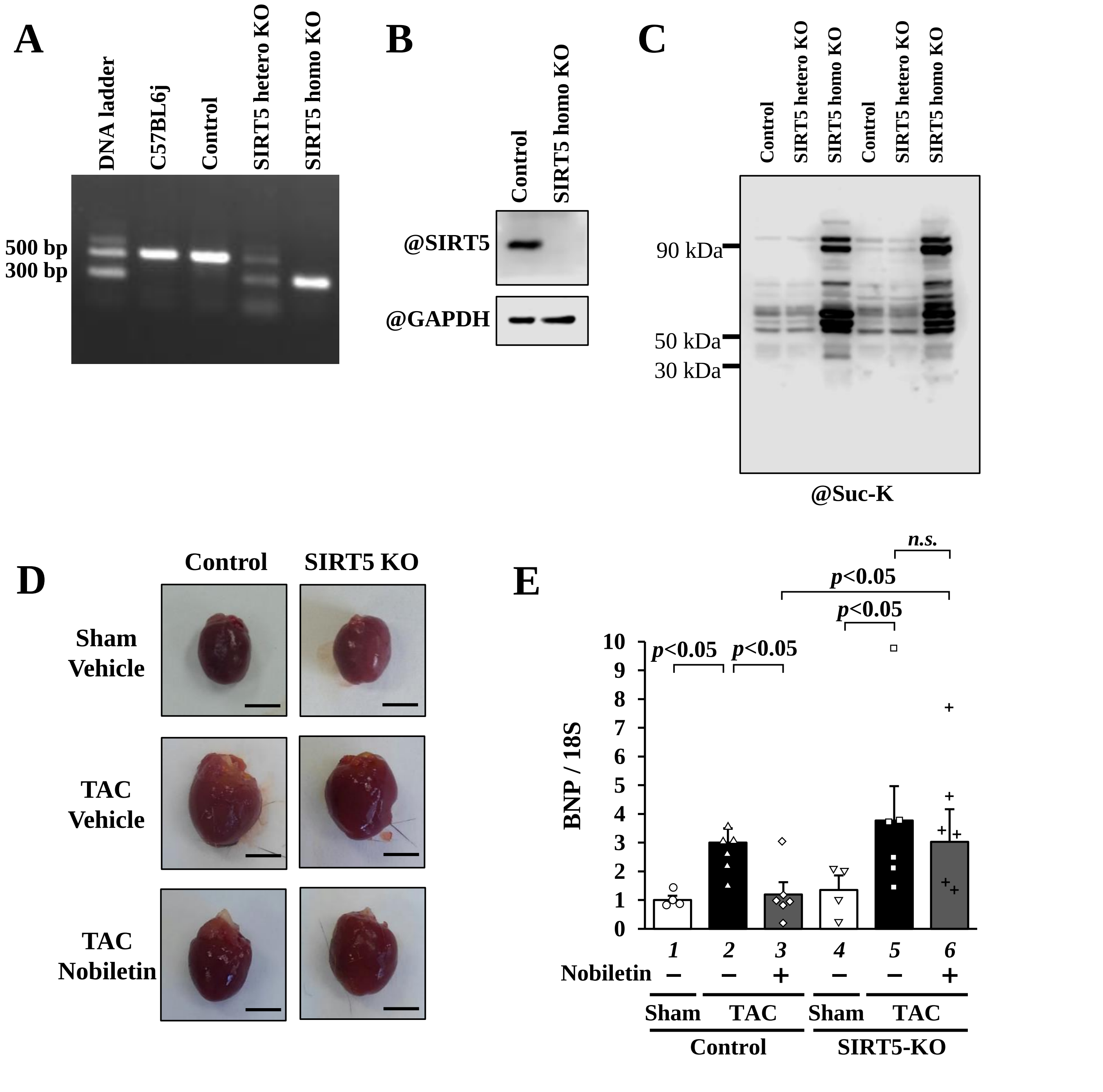


**Supplemental Figure .S10 The therapeutic potency of Nobiletin is abolished in SIRT5-KO mice**

**(A)** Genotyping of SIRT5-KO mice was detected in 1% agarose gel, a band of 300 bp was the knock-out allele, and a band of 500 bp was the WT allele. The extra bands in SIRT5 hetero KO mice indicate multiplex amplification. **(B)** Nuclear extracts from the hearts of SIRT5 homo KO and WT mice were subjected to western blotting using an anti-SIRT5 antibody. **(C)** Nuclear extracts from the hearts of SIRT5 homo KO, SIRT5 hetero KO, and WT mice were subjected to anti-succinylated lysine antibody. **(D)** The representative images of the whole from indicated groups at 6 weeks after surgery. Scale bar indicates 5 mm. **(E)** Total RNAs from each heart were subjected to quantified RT-PCR. The mRNA level of BNP was normalized by that of 18S. The values were shown as mean ± SEM.

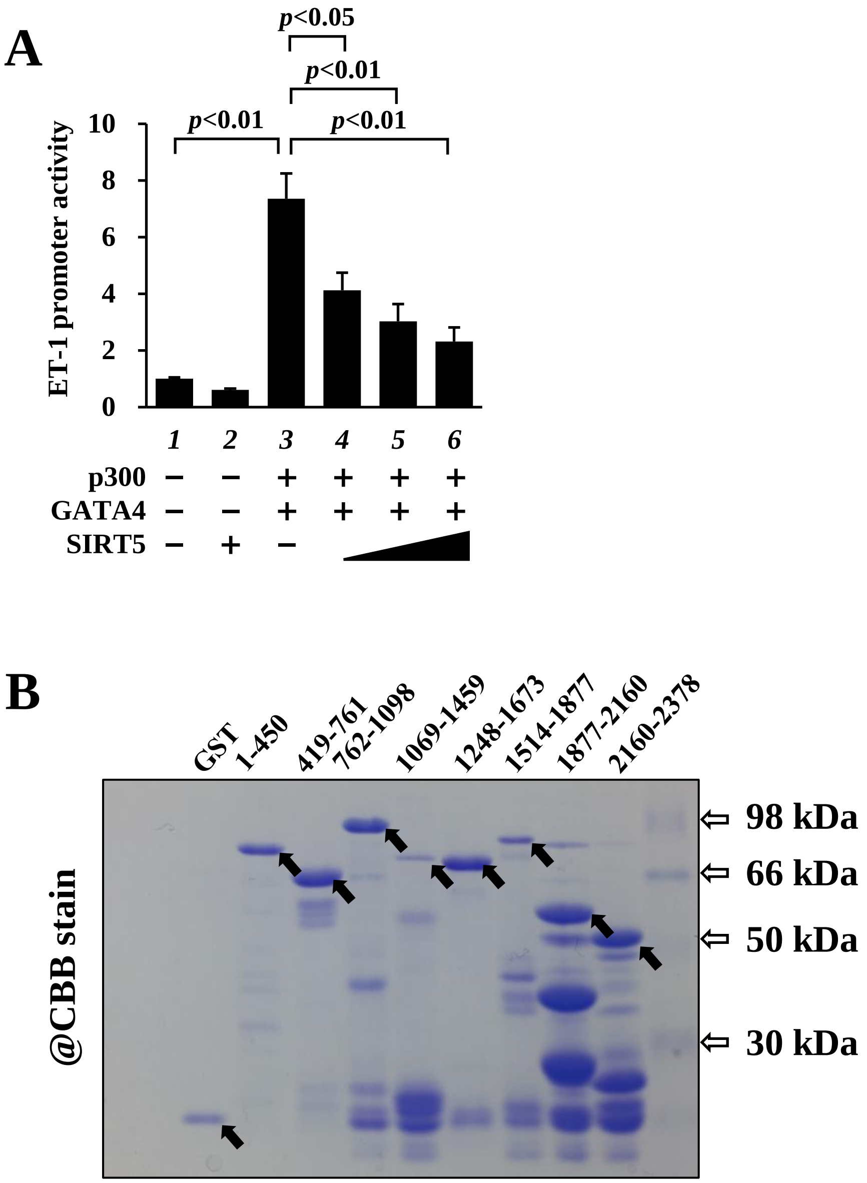


**Supplemental Figure. S11 SIRT5 reduces p300/GATA4-dependent promoter activation**

**(A)** HEK293T cells were transfected with pET-luc and pRL-SV40 in the presence or absence of pcDNA-GATA4, pcDNA-p300, and pcDNA-SIRT5. The relative promoter activities were shown as means ± SEM from three independent experiments, each carried out in duplicate. **(B)** Recombinant V5-SIRT5 was incubated with indicated GST-fused deletion mutants of p300 or GST protein as a control for 2 h. Precipitated GST-fused mutants and GST protein were detected by coomassie brilliant blue (CBB) staining. The arrows indicated GST proteins.

### Supplemental Figure S12

**A**

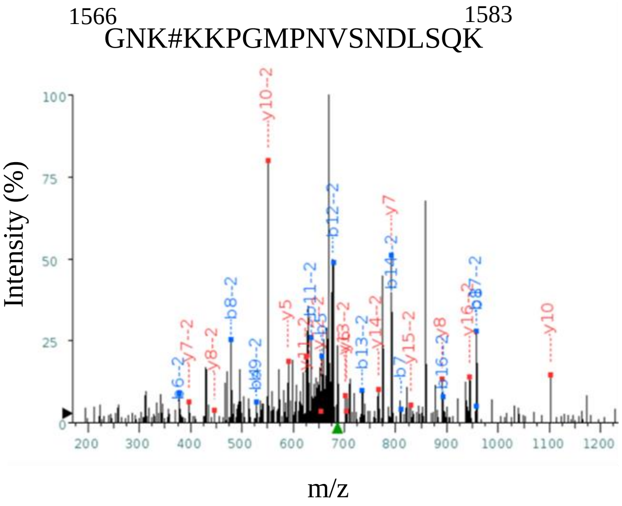


**B**

Sequence

GNK#KKPGM*PNVSNDLSQK

Predicted Fragmentation Pattern

+1

| Seq | # | b: Δ Error | b | y | y: Δ Error | +1 |
|---|---|---|---|---|---|---|
| G | 1 | --- | 58.029 | --- | --- | 18 |
| N | 2 | --- | 172.072 | 2001.007 | --- | 17 |
| K# | 3 | --- | 400.183 | 1886.964 | --- | 16 |
| K | 4 | -1545.264 | **528.278** | 1658.853 | --- | 15 |
| K | 5 | 559.389 | **656.373** | 1530.758 | --- | 14 |
| P | 6 | --- | 753.425 | 1402.663 | --- | 13 |
| G | 7 | 852.006 | **810.447** | 1305.610 | --- | 12 |
| M* | 8 | 237.772 | **957.482** | 1248.589 | --- | 11 |
| P | 9 | --- | 1054.535 | **1101.553** | -9.652 | 10 |
| N | 10 | --- | 1168.578 | 1004.501 | --- | 9 |
| V | 11 | --- | 1267.646 | **890.458** | 311.750 | 8 |
| S | 12 | --- | 1354.678 | **791.389** | 70.294 | 7 |
| N | 13 | --- | 1468.721 | **704.357** | 226.571 | 6 |
| D | 14 | --- | 1583.748 | **590.314** | 236.571 | 5 |
| L | 15 | --- | 1696.832 | 475.287 | --- | 4 |
| S | 16 | --- | 1783.864 | 362.203 | --- | 3 |
| Q | 17 | --- | 1911.923 | 275.171 | --- | 2 |
| K | 18 | --- | --- | 147.113 | --- | 1 |

**Supplemental Figure. S12 The lysine residues of p300 at 1568, 1569, and 1570 are candidates for SIRT5-mediated de-succinylation**

**(A)** Collision-induced dissociation mass spectrum of the human p300 derived succinylated peptide (1566-1583). # indicates a candidate of succinylated lysine residue. **(B)** Predicted fragmentation pattern of the human p300 derived succinylated peptide (1566-1583).

### Supplemental Figure S13

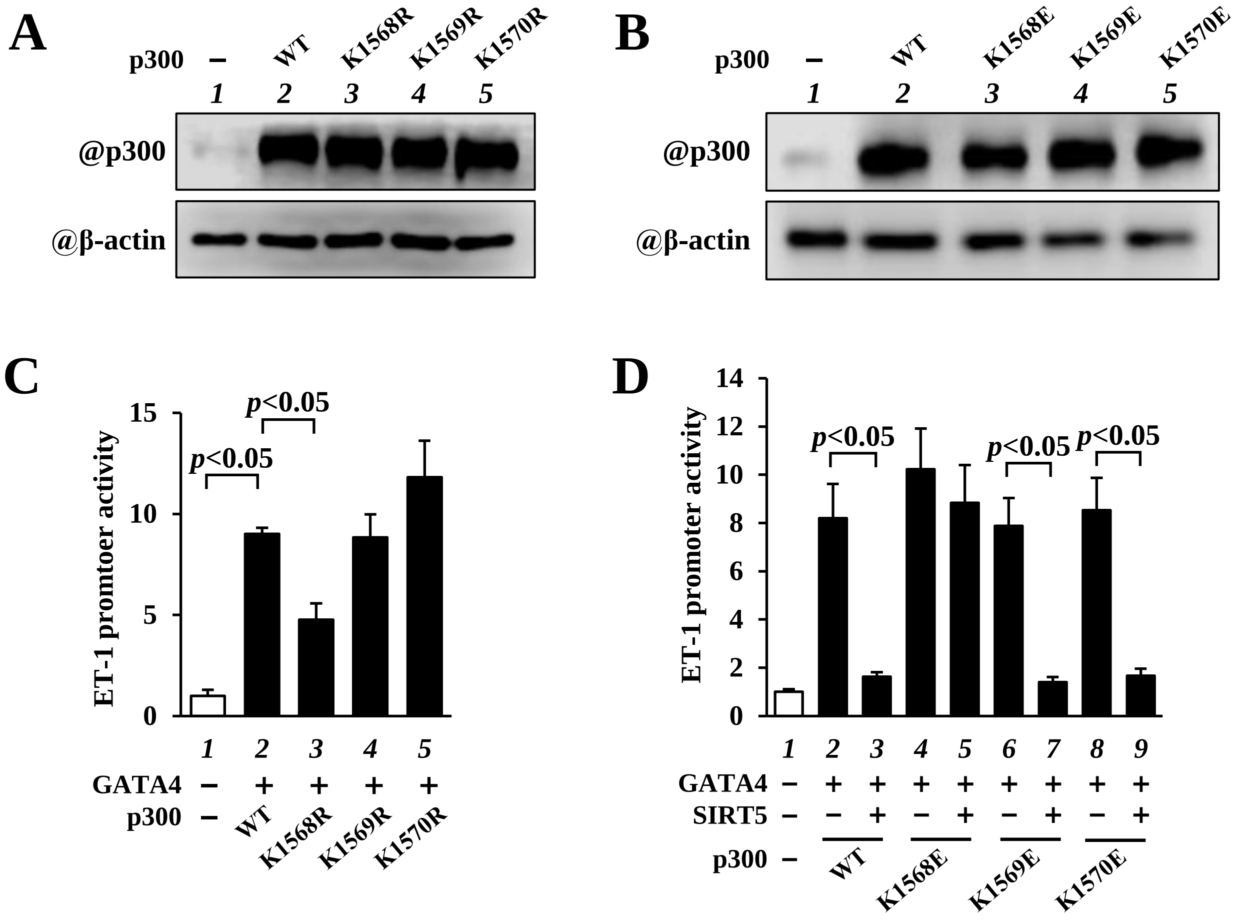


**Supplemental Figure. S13 The succinylation of a lysine residue of p300 at 1568 is required for p300-dependent hypertrophic responses**

HEK293T cells were transfected with expression vectors encoding p300WT or indicated p300 mutants. Nuclear extractions from these cells were subjected to western blotting. **(A)** The K-to-R-substituted p300 mutants. **(B)** The K-to-E-substituted p300 mutants. **(C and D)** HEK293T cells were transfected with pET-luc, pRL-SV40, indicated vectors, and p300 mutants. The relative promoter activities were shown as means ± SEM from three independent experiments, each carried out in duplicate.

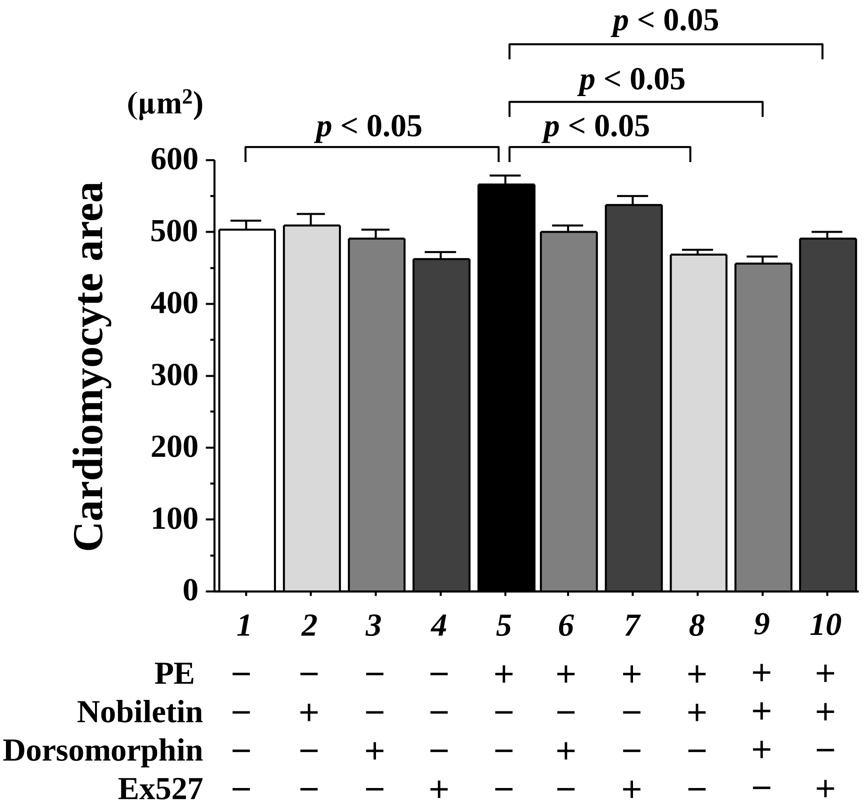


**Supplemental Figure. S14 AMPK/SIRT1 axis is not involved in the anti-hypertrophic effect of Nobiletin**

Cardiomyocytes were treated with Dorsomorphin, an inhibitor of adenosine monophosphate-activated protein kinase (AMPK), Ex527, an inhibitor of SIRT1, or DMSO as a control in the presence or absence of Nobiletin and stimulated with PE for 48 h. The surface area of fifty β-MHC-positive cardiomyocytes was automatically measured. The data were shown as means ± SEM from three independent experiments.
