## Supplemental table for "Nobiletin, a Polymethoxyflavonoid, Activates the Desuccinylase Activity of SIRT5 and Prevents the Development of Heart Failure"

### Supplemental Table S1

**The primer sequences in this study**

| Primer | | Sequence |
|---|---|---|
| Mouse ANF | Forward | 5’-TTCCTCGTCTTGGCCTTTTG-3’ |
| | Reverse | 5’-CCTCATCTTCTACCGGCATCTTC-3’ |
| Mouse BNP | Forward | 5’-AAGCTGCTGGAGCTGATAAGA-3’ |
| | Reverse | 5’-GTTACAGCCCAAACGACTGAC-3’ |
| Mouse 18S | Forward | 5’-CTTAGAGGGACAAGTGGCG-3’ |
| | Reverse | 5’-GGACATCTAAGGGCATCACA-3’ |
| Rat ANF | Forward | 5’-ATCACCAAGGGCTTCTTCCT-3’ |
| | Reverse | 5’-CCTCATCTTCTACCGGCATC-3’ |
| Rat BNP | Forward | 5’-ATCTGCCCTCTTGAAAAGCA-3’ |
| | Reverse | 5’-TCGAGCAGATTTGGCTGTTA-3’ |
| Rat 18S | Forward | 5’-GGTGCATGGCCGTTCTTA-3’ |
| | Reverse | 5’-TCGTTCGTTATCGGAATTAACC-3’ |
| Mouse SIRT5 | Forward | 5’-CCCAGAGCCAGAGACTCAAG-3’ |
| | Reverse | 5’-CAGAGGATGTTCCCACCACT-3’ |
| Rat SIRT5 | Forward | 5’-GTCATCACCCAGAACATTGA-3’ |
| | Reverse | 5’-ACGTGAGGTCGCAGCAAGCC-3’ |
| Genotyping SIRT5 OE | Forward | 5’-GTCATCACCCAGAACATCGA-3’ |
| | Reverse | 5’-ACGTGAGGTCGCAGCAAGCC-3’ |
| Genotyping SIRT5 KO | Forward1 | 5’-TCATTCTCAGTATTGTTTTGCC-3’ |
| | Forward2 | 5’-AGGAGGTGGCAAAGGTCTTGC-3’ |
| | Reverse | 5’-CTGAGGTAGAGTCTCTCATTG-3’ |

### Supplemental Table S2

#### The echocardiographic parameters at 8 weeks after TAC operation

| Operation | Treatment | n | LVEDD (mm) | LVEDS (mm) | LVEF (%) | HR (bpm) | BW (g) |
|---|---|---|---|---|---|---|---|
| Sham | Vehicle | 5 | 2.47±0.08 | 1.08±0.06 | 93.0±0.9 | 447±19 | 26.3±0.7 |
| Sham | Nobiletin 20 mg/kg | 5 | 2.46±0.10 | 1.00±0.10 | 92.3±1.3 | 498±14 | 25.1±1.0 |
| TAC | Vehicle | 8 | 2.77±0.17 | 1.68±0.17* | 77.0±3.1* | 476±14 | 26.1±0.3 |
| TAC | Nobiletin 20 mg/kg | 7 | 2.40±0.16 | 1.04±0.20## | 88.5±5.9# | 455±13 | 25.5±0.4 |

*: $p < 0.05$, **: $p < 0.01$ *vs* sham vehicle
#: $p < 0.05$, ##: $p < 0.01$ *vs* TAC vehicle

LVEDD: left ventricular end-diastolic dimension, LVEDS: left ventricular end-systolic dimension, LVEF: left ventricular ejection fraction, HR: heart rate, SBP: systolic blood pressure, DBP: diastolic blood pressure, BW: body weight

### Supplemental Table S3

#### The echocardiographic and hemodynamic parameters at 1 weeks after post-MI

| Operation | Treatment | n | LVEDD (mm) | LVEDS (mm) | LVPWT (mm) | FS (%) | LVEF (%) | HR (bpm) | BW (g) |
|---|---|---|---|---|---|---|---|---|---|
| Sham | Vehicle | 5 | 7.39±0.24 | 3.22±0.28 | 1.09±0.24 | 56.7±2.6 | 91.6±2.6 | 386±20 | 318±4 |
| MI | Vehicle | 5 | 8.98±0.11* | 6.51±0.12* | 1.31±0.17 | 27.5±1.3* | 61.7±1.5# | 394±14 | 324±5 |
| MI | Nobiletin 1 mg/kg | 5 | 9.04±0.32* | 6.53±0.30* | 1.32±0.10 | 27.8±1.5* | 62.2±1.5# | 387±26 | 325±4 |
| MI | Nobiletin 20 mg/kg | 5 | 8.96±0.24* | 6.49±0.31* | 1.36±0.04 | 27.8±1.7* | 62.0±1.7# | 400±22 | 333±15 |

*:$p < 0.05$, **: $p < 0.01$ *vs* sham vehicle
#:$p < 0.05$, ##: $p < 0.01$ *vs* MI vehicle

LVEDD: left ventricular end-diastolic dimension, LVEDS: left ventricular end-systolic dimension, LVPWT: left ventricular posterior wall thickness, FS: fractional shortening, LVEF: left ventricular ejection fraction, HR: heart rate, SBP: systolic blood pressure, DBP: diastolic blood pressure, BW: body weight

### Supplemental Table S4

#### The echocardiographic and hemodynamic parameters at 7 weeks after post-MI

| Operation | Treatment | n | LVEDD (mm) | LVEDS (mm) | LVEF (%) | HR (bpm) | SBP (mmHg) | DBP (mmHg) | BW (g) | Infarct size (%) |
|---|---|---|---|---|---|---|---|---|---|---|
| Sham | Vehicle | 5 | 8.34±0.17 | 3.95±0.14 | 89.4±0.5 | 351±15 | 108±3 | 84±4 | 482±8 | - |
| MI | Vehicle | 5 | 10.31±0.19* | 8.53±0.18* | 43.4±1.3* | 386±13 | 124±4* | 91±3* | 456±9 | 31.2±4.7 |
| MI | Nobiletin 1 mg/kg | 5 | 10.53±0.48* | 8.47±0.41* | 47.4±1.5* | 421±22 | 118±3* | 89±2* | 452±9 | 30.1±3.3 |
| MI | Nobiletin 20 mg/kg | 5 | 11.22±0.15* | 8.32±0.24* | 59.3±1.7*# | 376±17 | 124±3* | 101±3* | 475±10 | 31.4±2.4 |

*: $p < 0.05$, **: $p < 0.01$ *vs* sham vehicle
#: $p < 0.05$, ##: $p < 0.01$ *vs* MI vehicle

LVEDD: left ventricular end-diastolic dimension, LVEDS: left ventricular end-systolic dimension, LVEF: left ventricular ejection fraction, HR : heart rate, SBP: systolic blood pressure, DBP: diastolic blood pressure, BW: body weight

### Supplemental Table S5

#### The echocardiographic parameters at 8 weeks after TAC operation in SIRT-TG mice

| Mice | Operation | n | LVEDD (mm) | LVEDS (mm) | LVEF (%) | HR (bpm) | BW (g) |
|---|---|---|---|---|---|---|---|
| WT mice | Sham | 5 | 2.78±0.17 | 1.24±0.09 | 90.5±1.0 | 445±28 | 28.5±0.4 |
| SIRT5-TG mice | Sham | 5 | 2.83±0.19 | 1.16±0.09 | 92.5±0.85 | 439±22 | 28.9±1.0 |
| WT mice | TAC | 7 | 3.36±0.20* | 2.21±0.26** | 67.2±5.0* | 477±7 | 28.3±0.5 |
| SIRT5-TG mice | TAC | 7 | 2.44±0.17# | 1.05±0.06## | 56.6±1.2# | 440±21 | 27.2±0.6 |

*: $p < 0.05$, **: $p < 0.01$ *vs* WT mice with sham
#: $p < 0.05$, ##: $p < 0.01$ *vs* WT mice with TAC

LVEDD: left ventricular end-diastolic dimension, LVEDS: left ventricular end-systolic dimension, LVEF: left ventricular ejection fraction, HR: heart rate, SBP: systolic blood pressure, DBP: diastolic blood pressure, BW: body weight

### Supplemental Table S6

**The echocardiographic parameters at 6 weeks after TAC operation in SIRT5-KO mice**

| Mice | Operation | Treatment | n | LVEDD (mm) | LVEDS (mm) | LVEF (%) | HR (bpm) | BW (g) |
|---|---|---|---|---|---|---|---|---|
| WT mice | Sham | Vehicle | 5 | 2.97±0.08 | 1.31±0.03 | 90.9±0.5 | 430±16 | 26.7±0.7 |
| WT mice | TAC | Vehicle | 7 | 3.19±0.06 | 1.85±0.07* | 79.2±1.6** | 458±19 | 25.6±0.9 |
| WT mice | TAC | Nobiletin 20 mg/kg | 7 | 3.32±0.16 | 1.66±0.10# | 86.6±0.7# | 450±26 | 25.4±1.0 |
| SIRT5-KO mice | Sham | Vehicle | 5 | 3.25±0.14 | 1.57±0.13 | 87.9±1.5 | 459±24 | 26.8±0.9 |
| SIRT5-KO mice | TAC | Vehicle | 7 | 3.44±0.16 | 2.26±0.14**†† | 70.3±2.1**#†† | 420±22 | 24.2±0.8 |
| SIRT5-KO mice | TAC | Nobiletin 20 mg/kg | 7 | 3.26±0.11 | 2.15±0.12**† | 70.0±3.0**#†† | 460±25 | 24.6±0.7 |

*: $p < 0.05$, **: $p < 0.01$ *vs* WT mice with sham
#: $p < 0.05$, ##: $p < 0.01$ *vs* WT mice with TAC
†: $p < 0.05$, ††: $p < 0.01$ *vs* KO mice with sham

LVEDD: left ventricular end-diastolic dimension, LVEDS: left ventricular end-systolic dimension, LVEF: left ventricular ejection fraction, HR: heart rate, SBP: systolic blood pressure, DBP: diastolic blood pressure, BW: body weight
